# A newly-identified mini-hairpin shaped nascent peptide blocks translation termination by a novel mechanism

**DOI:** 10.1101/2024.05.31.596752

**Authors:** Yushin Ando, Akinao Kobo, Tatsuya Niwa, Ayako Yamakawa, Suzuna Konoma, Yuki Kobayashi, Osamu Nureki, Hideki Taguchi, Yuzuru Itoh, Yuhei Chadani

**Affiliations:** Department of Biological Sciences, Graduate School of Science, The University of Tokyo, Tokyo 113-0033, Japan; School of Life Science and Technology, Tokyo Institute of Technology, Yokohama 226-8501, Japan; Cell Biology Center, Institute of Innovative Research, Tokyo Institute of Technology, Yokohama 226-8501, Japan; Faculty of Environmental, Life, Natural Science and Technology, Okayama University, Okayama 700-8530, Japan

**Keywords:** nascent polypeptide, ribosome arrest peptide, cryo-EM, translation termination, release factor (RF), stop codon read-through, ribosome stalling, peptidyl-tRNA, *Escherichia coli*

## Abstract

Protein synthesis by ribosomes not only produces functional proteins but also serves diverse functions depending on the coding amino acid sequences. Certain nascent peptides interact with the ribosome exit tunnel to arrest translation and modulate the expression of downstream genes or themselves. However, a comprehensive understanding of the mechanisms of such ribosome stalling and its regulation remains elusive. In this study, we systematically screened for unidentified ribosome arrest peptides through phenotypic evaluation, proteomics, and MS analyses, leading to the discovery of novel arrest peptides PepNL and NanCL in *E. coli*. Our cryo-EM study on PepNL revealed a unique arrest mechanism, in which the N-terminus of PepNL folds back towards the tunnel entrance to prevent the catalytic GGQ motif of release factor from accessing the peptidyl transferase center, causing translation arrest at the UGA stop codon. Furthermore, unlike other sensory arrest peptides that require an arrest inducer, PepNL uses tryptophan as an arrest releaser, where Trp-tRNA reads through the stop codon. Our findings illuminate the mechanism and regulatory framework of nascent peptide-induced translation arrest, paving the way for exploring regulatory nascent peptides.

## Introduction

Translation is a crucial process that converts nucleotide sequences into amino acid sequences, facilitating the expression of a diverse array of proteins with various functions inside the cell. This complex process is orchestrated by the ribosome, a universal protein factory found in all forms of life on Earth. In bacteria, ribosomes assemble an initiation complex upon encountering the Shine-Dalgarno sequence in mRNA upstream of the initiation codon (**1**). Subsequently, ribosomes catalyze the transfer of the growing peptide chain from aminoacyl/peptidyl-tRNA at the P-site to aminoacyl-tRNA at the A-site (**2**). Upon encountering a stop codon, peptide release factors (RF) recognize the stop codon to trigger the hydrolysis of the ester bond between the nascent peptide and tRNA (**3–8**). This event leads to the dissociation of the polypeptide chain and terminates translation.

While ribosomes were once considered robust protein factories capable of translating any mRNA sequence, it is now evident that this is not always the case. One such case is that the nascent polypeptide chain (nascent peptide) within the ribosome tunnel causes translational difficulties (**9–11**). The nascent peptide exit tunnel (NPET) is primarily composed of negatively charged rRNA and contains a constriction site formed by ribosomal proteins (**12**). Certain amino acid sequences in the nascent peptide have been shown to closely interact with the complex tunnel structures, causing the elongation stalling termed “translation arrest” (**13, 14**). Although translation arrest appears to be disadvantageous and might be eliminated during evolution, there are conserved translation arrests among organisms, which may utilize it as part of their survival strategies (**15–19**).

The majority of ribosome arrest peptides (RAPs) discovered so far are encoded within upstream ORFs (uORFs) (**9, 10**). The occurrence of the translation arrest often depends on environmental changes inside and outside the cell to regulate the expression of downstream genes. For example, *Escherichia coli* TnaC induces translation arrest only when the intracellular tryptophan concentration is high (**20**). This is because excessive tryptophan enters the ribosome tunnel, to alter the structure of the peptidyl transferase center (PTC) structure through interactions with the nascent peptide and the tunnel, inhibiting the translation termination (**21, 22**). In this way, TnaC regulates the expression of tryptophanase (TnaA) in a manner dependent on intracellular tryptophan as an “arrest inducer”. RAPs that control the expression of downstream genes like TnaC are widely found from prokaryotes to eukaryotes and are considered useful and universal gene expression regulation mechanisms in life (**9–11**). The search for such unidentified RAPs has been advanced through sequence and gene structure conservation analyses (**23–26**), as well as large-scale translatome analyses using ribosome profiling (**27, 28**). However, the overall landscape of nascent peptide-dependent translation regulation remains elusive.

In addition to the diversity of the sequences that cause translation arrest, the molecular mechanisms that cause arrest are also diverse. Recent cryo-electron microscopy (cryo-EM) structural analyses have significantly advanced our understanding of the molecular details of translation arrest. Several recent reports have revealed intricate interactions between RAPs and the ribosome tunnel at resolutions approaching 2.0 Å (**21, 22, 29–36**). Structural studies of arrest peptides such as TnaC and SpeFL revealed the metabolite-sensing and release factor-inhibiting mechanisms. However, structural analyses of ribosomes arrested by RAPs are currently limited to typical examples, and further case studies are needed to systematically understand the mechanism of translation arrest.

This study comprehensively examined whether small ORFs newly identified from ribosome profiling exhibit RAP activity. Through phenotype characterization, proteomic analysis, and mass spectrometry (MS) analyses of nascent peptides, *E. coli* PepNL (14 aa) and NanCL (14 aa) were demonstrated to induce translation arrest at their stop codons. Focusing on PepNL, our cryo-EM analysis revealed an unusual structure where the nascent peptide of PepNL folds back towards the entrance of the ribosome tunnel, contrary to the usual orientation. This distinctive conformation distorts the structure of the subsequent nascent peptide, leading to a steric clash with the GGQ motif of RF2, thereby inhibiting translation termination. Unlike previously studied RAPs such as TnaC, PepNL arrested the ribosome solely based on its amino acid sequence, without requiring any arrest inducer. This study not only unveiled a novel mode of translation arrest but also provided insights into the associated mechanism for the release of PepNL-dependent arrest.

## Results

### Screening of novel ribosome arrest peptides by over-expression phenotype

Previously, it was reported that over-expression of TnaC, a tryptophan-dependent ribosome arrest peptide (RAP), impedes cell growth due to the depletion of tRNA^Pro^ captured within the arrested ribosome (**Fig. 1a**) (**37**). To identify novel RAPs, we investigated whether the over-expression of RAPs other than TnaC would inhibit cell growth, as observed with TnaC in nutrient-rich media. Notably, over-expression of established RAPs in *E. coli* cells growing on LB agar, such as SecM (**13**) and TnaC (**14**), resulted in significant growth inhibition (**Fig. 1b**). In contrast, SpeFL, which requires a sufficient supply of ornithine for arrest induction (**Fig. 1b**) (**33**), or the TnaC variant harboring the arrest-attenuating P24A mutation, failed to induce growth inhibition (**Supplementary Fig. 1a**). MgtL, another type of regulatory nascent peptide that triggers premature translation abortion, had a negligible effect on cell growth (**Fig. 1b**) (**38**). Based on these results, we inferred that the cytotoxicity upon over-expression could serve as one of the indicators of RAP activity.

**Figure 1.**
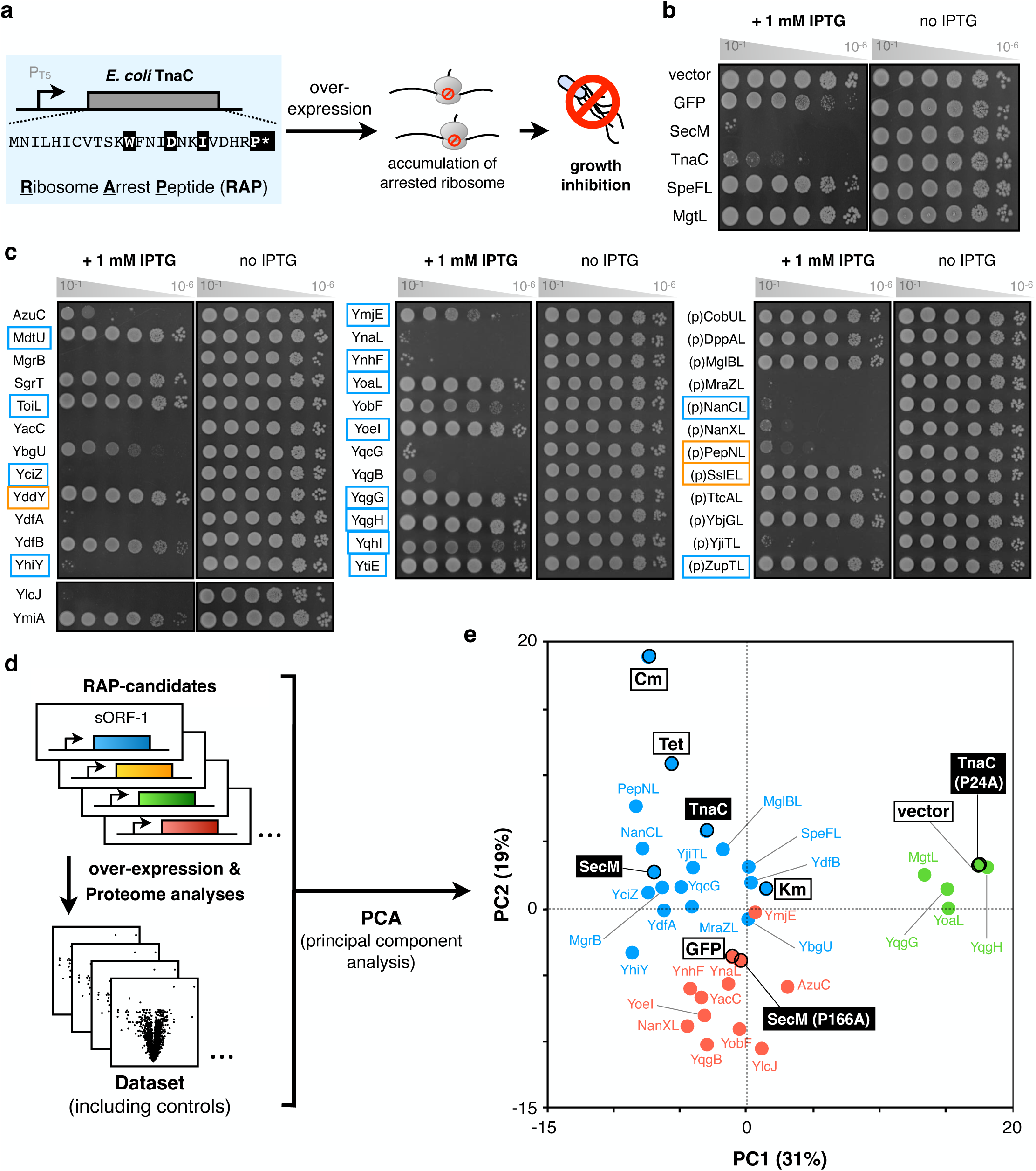
Screening of novel ribosome arrest peptides by over-expression phenotype. **a**. Working hypothesis for screening novel ribosome arrest peptide. **b and c**. Serial dilution assay for evaluating the cytotoxicity when established (**b**) or recently identified (**c**) sORFs were over-expressed in *E. coli* cells. **d**. A schematic illustration of a comparative quantitative proteomic analysis. Each sORF was over-expressed in *E. coli* cells, and all the samples were applied to LC-MS/MS-based proteome analysis. The proteomic changes by the over-expression were quantified as the fold changes against the cells harboring an empty vector. **e**. PCA score plot of PC1 and PC2. The numbers in the x and y-axis labels represent the proportion of the variances of PC1 and PC2, respectively. Vector means the data from fold changes of another vector control against a vector control. Colors (green, blue, and red) correspond to the clustering by a hierarchical clustering analysis (**Supplementary Fig. 1i**).

In this study, we analyzed a total of 38 candidates, including 26 recently annotated small open reading frames (sORFs) (**27, 28, 39–41**) and 12 putative sORFs identified by using the GWIPS-viz browser (**Supplementary Fig. 1b-d)** (**42**). Upon over-expression of these 38 candidates in *E. coli* cells, 18 sORFs induced growth inhibition (**Fig. 1c**). The presence or absence of cytotoxicity did not correlate with their regulatory effects on downstream gene expression (**Fig. 1c, colored frame, Supplementary Fig. 1e**).

Subsequently, we investigated whether the examined sORFs induce stress responses due to excessive translation arrest. It is anticipated that over-expression of TnaC would inhibit translation elongation by tRNA^Pro^ depletion. In such a situation, the expression of cold shock proteins (CSPs) is expected to increase, as observed in *E. coli* cells exposed to sublethal concentrations of chloramphenicol (Cm) or tetracycline (Tet), which globally inhibit translational elongation (**43–46**). Based on this hypothesis, we conducted a comparative and quantitative proteomic analysis for *E. coli* cells over-expressing each of the sORFs, and analyzed their proteomic landscape rearrangement (**Fig. 1d**). The obtained dataset (**Supplementary Dataset 1 and 2**) was subjected to PCA (principal component analysis), and a plot illustrating their similarity of proteomic rearrangement is depicted in **Fig. 1e**. Parallel clustering analysis categorized our datasets into three clusters: I. induction of CSPs expression (blue circles), II. induction of HSPs expression (red circles), III. limited variation (green circles) (**Fig. 1e, Supplementary Fig. 1f-i**). Over-expression of RAPs such as TnaC and SecM, as well as treatment with translation inhibitors like Cm, induced the expression of CSPs, such as CspA, DeaD, and InfC (**Supplementary Fig. 1h**) (**47–50**). These datasets were grouped in the same cluster, indicating that inhibition of the translation elongation often induces the expression of CSPs (**Fig. 1e blue circles, Supplementary Fig. 1i**). A similar “cold shock”-like response was observed for 12 sORF candidates, indicating the possible occurrence of translation arrest during their expression. On the other hand, several sORFs such as YqgB induced the expression of heat shock proteins (HSPs) that strongly respond to aggregation, including IbpA and ClpB (**Fig. 1e red circles, Supplementary Fig. 1h and 1i**) (**43, 51, 52**). Over-expression of these sORFs, unlike TnaC, might inhibit cell growth due to aggregation of the expressed proteins.

### LC-MS/MS measurements of the peptidyl-tRNA molecules

We further evaluated the RAP activity of candidate sORFs by LC-MS/MS analysis of the peptidyl-tRNA intermediates that accumulate due to translation arrest. After the over-expression of sORFs in *E. coli* cells, total RNA was extracted and purified by silica adsorption. Peptide fragments ester-bonded to tRNA were then subjected to alkaline hydrolysis, peptidase digestion, and LC-MS/MS analysis (**Fig. 2a**) (**53, 54**). Identification of the MS2 spectra of the ^19th^ K / IVDHRP^25th^ fragment of TnaC (**Fig. 2b**) or ^149th^K / GSTPVWISQAQGIRAG^165th^ fragment of SecM (**Fig. 2c**), which coincides with the established translation arrest site, demonstrates the effectiveness of this method. Additionally, the identification of Tris-adducted peptide fragments at the C-terminus, which occurs upon alkaline hydrolysis of peptidyl-tRNA, further supports the RAP activity of TnaC and SecM (**Supplementary Fig. 2a and 2b**).

**Figure 2.**
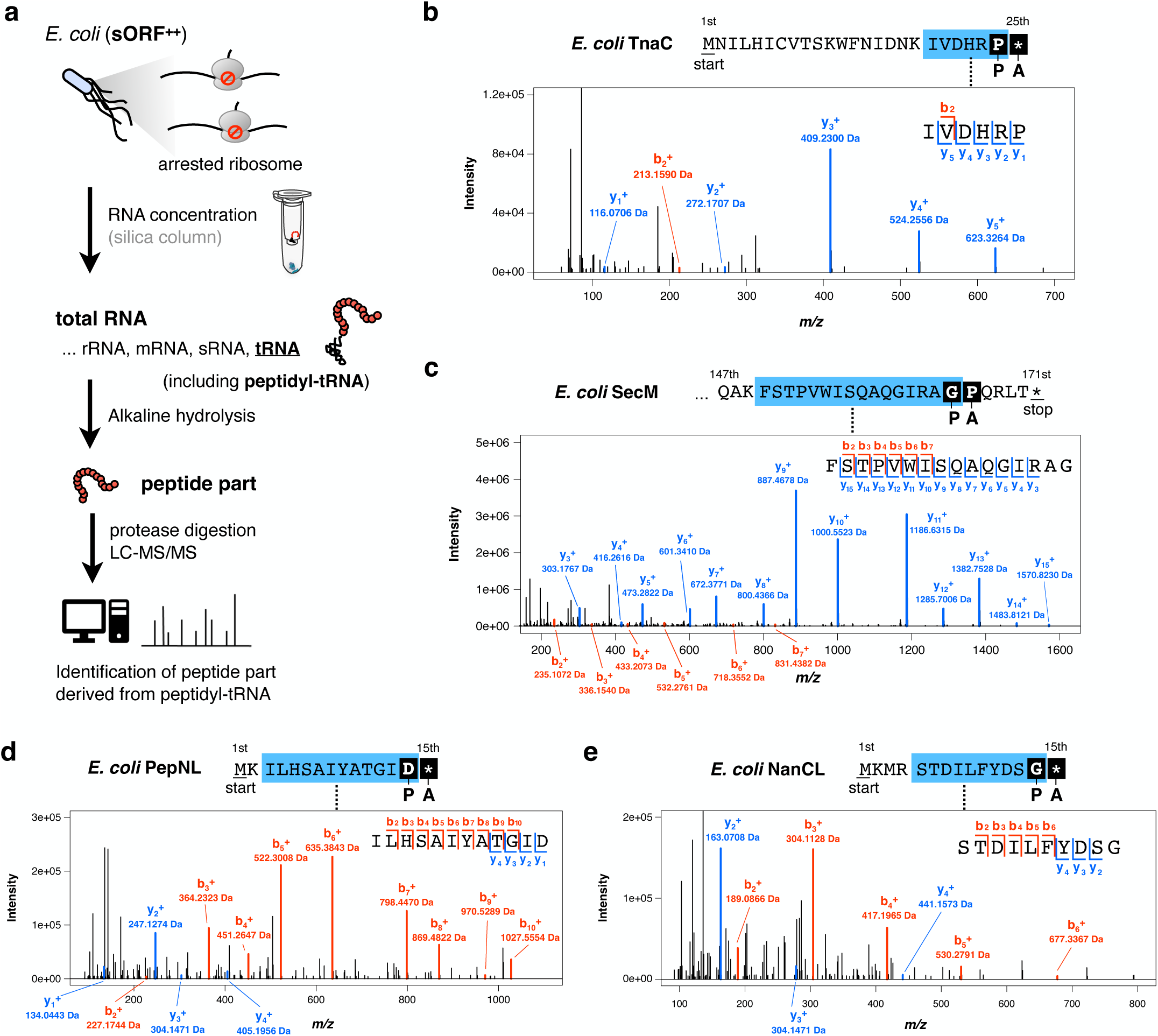
Identification of the ribosome-arresting peptidyl-tRNA by LC-MS/MS. **a.** A schematic illustration of the experimental procedure for the identification of peptidyl-tRNA-derived peptides. Each candidate sORF was over-expressed in *E. coli* cells, and the total RNA fraction in the cell lysate was concentrated with a silica column. Then, the fraction was hydrolyzed by alkaline treatment, and the obtained peptides were identified by protease digetion and subsequent LC-MS/MS. **b-e.** MS/MS spectra of peptides derived from peptidyl-tRNAs. Amino acid sequences above the graphs represent the positions of the detected peptides (blue areas indicate the detected peptide). P and A below the amino acid sequence represent the plausible positions of the P- and A-sites in the ribosome during translation arrest, respectively. In the graphs, the peaks of the b- and y-fragment ions are shown in red and blue, respectively.

The LC-MS/MS analysis of peptidyl-tRNA was extended to include sORF candidates, resulting in the identification of peptide fragments indicative of stalling at various sites (**Supplementary Table 1**). Many of the identified peptide fragments were inconsistent with the distribution of the ribosome on the mRNA in the ribosome profiling. In contrast, the MS2 spectra indicating translation arrest at stop codons of the sORF candidates *pepNL* and *nanCL* were in line with the profiling results (**Fig. 2d and 2e, Supplementary Fig. 1b**). In addition, Tris-adducted peptide fragments at the C-terminus were detected for these two sORFs, as in the case of TnaC and SecM (**Supplementary Fig. 2c and 2d**). Based on these LC-MS/MS results and the growth inhibition results shown in **Fig. 1**, we concluded that *pepNL* and *nanCL* encode novel RAPs that induce translation arrest at stop codons.

### Structure of the PepNL-arrested ribosome

In this study, we focused on PepNL, which induced a particularly robust translation stress response (**Fig. 1e**). PepNL, consisting of 14 amino acids, is encoded upstream of the aminopeptidase *pepN* gene. The frameshift mutation abolished the cytotoxicity (**Fig. 3a**) and the accumulation of peptidyl-tRNA (**Fig. 3b**), indicating that PepNL arrests translation through its context of the amino acid sequence. Moreover, the frameshift mutation increased the expression of PepN, underscoring the regulatory role of *pepNL* in a translation arrest-dependent manner (**Supplementary Fig. 1e and 3a**).

**Figure 3.**
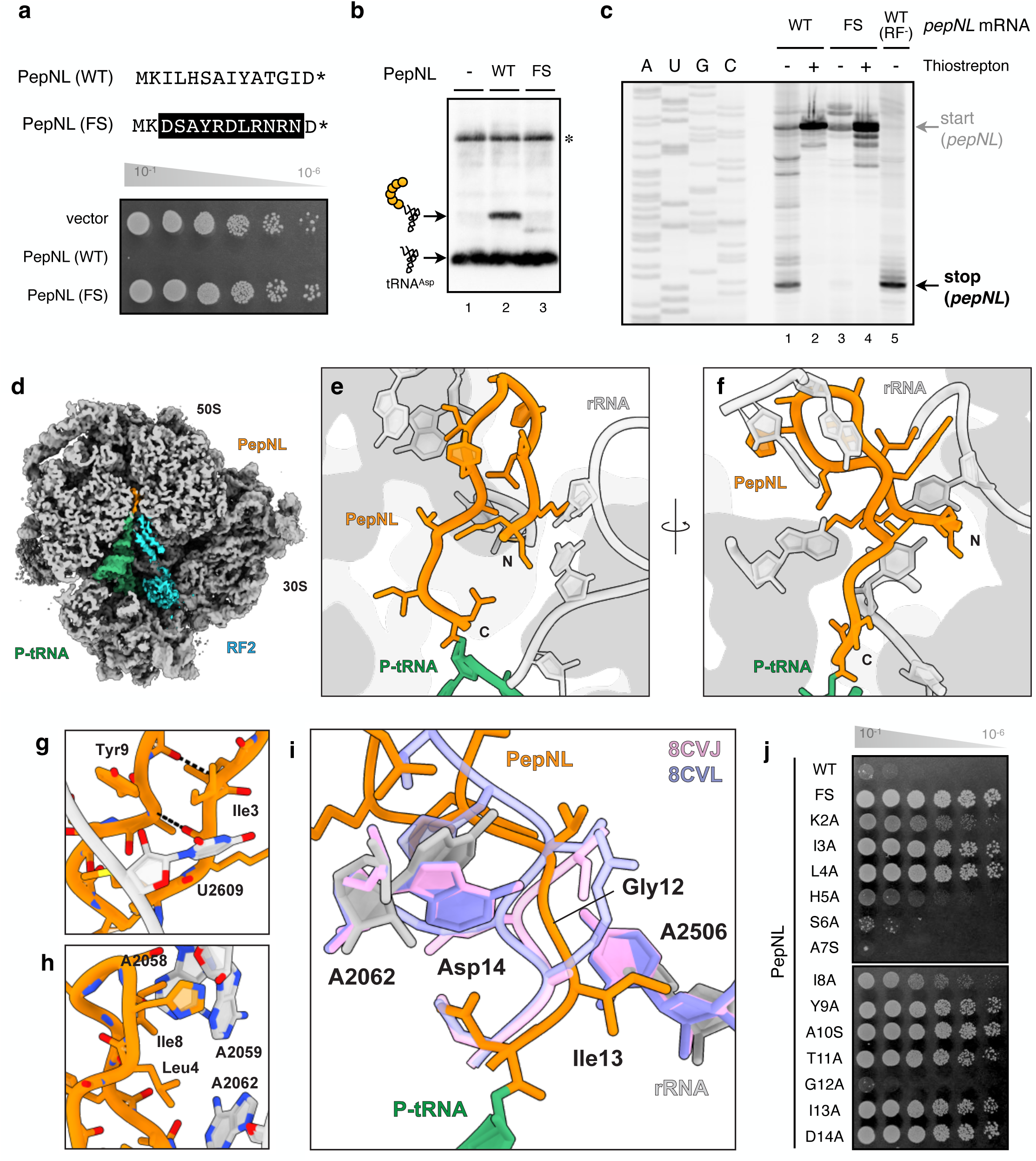
PepNL nascent peptide turns back toward the entrance of the ribosome tunnel. **a.** Serial dilution assay to assess the cytotoxicity upon over-expression of wild-type PepNL (WT) or its frameshift mutant (FS) in *E. coli* cells. **b.** The PepNL-tRNA accumulated in *E. coli* cells expressing the indicated *pepNL* variants was detected by northern blotting using anti-tRNA^Asp^ probe. An asterisk indicates the unprocessed tRNA transcript. **c.** The wild-type or frameshifted *pepNL* mRNA was translated by PURE*frex* in the absence of tryptophan, and the PepNL-arrested ribosome was visualized by toeprint analysis. Thiostrepton, which inhibits translation elongation was pre-included where indicated. The *pepNL* mRNA was translated in the absence of release factors (RF^-^) to prepare the ribosomes stalled at the stop codon for the position marker (lane 5). **d**, Overall cross section of the cryo-EM density map at the peptide exit tunnel of 70S ribosome (gray), showing P-tRNA (green), RF2 (cyan), and PepNL peptide (orange). **e, f**, PepNL peptide and interacting 23S rRNA nucleotides in the ribosome exit tunnel. **g, h**, Close-up views of the intermolecular interactions within PepNL peptide (**g**) and intramolecular interactions between PepNL peptide and 23S rRNA nucleotides (**h**). **i**, Structure comparison with non-arrested states (PDB: 8CVJ, 8CVL). The nascent peptides and A2062 and A2506 nucleotides at PTC are shown. **j**. Serial dilution assay to assess the cytotoxicity upon over-expression of wild-type PepNL (WT) or its variants carrying the indicated amino acid substitution in *E. coli* cells.

To elucidate the molecular mechanism of PepNL-dependent translation arrest, we analyzed the structure of the ribosome arrested by the PepNL nascent peptide. We translated *pepNL* mRNA using the reconstituted cell-free translation system (PURE system: PURE*frex* v1.0) (**55**) deprived of tryptophan, which will be discussed later (**Fig. 5**). Toeprint analysis confirmed the formation of ribosomes stalled at the stop codon of PepNL (**Fig. 3c, Supplementary Fig. 3b**). To avoid dissociation or release of the stalled ribosomes, we directly applied the *in vitro* translation mixture to the cryo-EM grids without purification and performed cryo-EM data collection. The contrast of the ribosomes on the grid was sufficient for the cryo-EM analysis even without purification, resulting in the 3D reconstruction of ribosomes. We obtained four states with different components in the A and P-sites (P-tRNA only: 31%, RF2 and P-tRNA: 25%, EF-Tu•tRNA and P-tRNA: 23%, and empty: 21%, **Supplementary Fig. 4**). We further analyzed the particles in the RF2 and P-tRNA bound state and P-tRNA bound state, and determined their structure at the resolution of 2.9 Å (**Fig. 3d**, **Supplementary Fig. 5**). In the determined structure with RF2, the arrested 70S ribosome bound with the mRNA, RF2 at the A-site, and tRNA^Asp^ carrying the nascent PepNL peptide at the P-site.

All 14 amino acid residues of PepNL were successfully traced in the density, indicating that the PepNL nascent peptide forms a stable conformation in the exit tunnel. The PepNL peptide forms a unique hairpin conformation with residues from Lys2 to Ala10, directing the N-terminal residues back to the PTC, not reaching the constriction site of the exit tunnel (**Fig. 3e and 3f**). This distinctive structure is stabilized by intramolecular interactions, including a hydrophobic interaction between Ile3 and Tyr9 (**Fig. 3g**), as well as β-sheet-like main-chain interactions involving Lys2 with Ala10 and Leu4 with Ile8, respectively (**Fig. 3h**). Moreover, intermolecular hydrophobic interactions between PepNL (Ile3, Leu4, Ile8, and Tyr9) and 23S rRNA (U2609, A2062, A2058, and A2059) also contribute to supporting the structure (**Fig. 3g and 3h**). While the N-terminal portion of PepNL exhibits such compact packing conformation, the C-terminal part is distorted, illustrated by a large main-chain shift as compared to the non-arrested structure (**Fig. 3i**) (PDB: 8CVJ and 8CVL, **reference-56**). The distortion of PepNL further interrupts the hydrogen-bond network between rRNA (A2506 and A2062) and the nascent peptide observed in the non-arrested structure, resulting in the conformational rearrangement of A2506 and A2062 residues of 23S rRNA (**Fig. 3i**). These structural observations were supported by the fact that the arrest activity of PepNL was abolished by alanine substitutions of Ile3, Leu4, Ile8, and Tyr9, which are involved in the formation of the β-hairpin loop (**Fig. 3j, Supplementary Fig. 3c**).

### RF2 undergoes a conformational rearrangement

We next focused on the structural rearrangement of RF2. In canonical translation termination, the recognition of the stop codon by domain II of RF2 triggers the positional extension of domain III, inserting the ^250th^GGQ^252nd^ catalytic motif into the PTC to hydrolyze the ester bond of the peptidyl-tRNA (**Fig. 4a**) (**5–8**). Within the PepNL-arrested ribosome, the domain II of RF2 properly recognizes the UGA stop codon, and the extension of domain III is also observed (**Fig. 4a**). However, we observed a significant difference in the conformation of the apical loop (residues 246-257), when we compared our structure with the canonical termination complex (PDB: 6C5L, **reference-57**) (**Fig. 4b**). In the canonical structure, the methylated Gln252 of the GGQ motif enters the narrow pocket formed by A2451, C2452, and U2506 of rRNA (**Fig. 4b and 4f**). However, in the arrested structure, Gln252 undergoes a 16 Å relocation, as measured between the Nε2 atoms (**Fig. 4b**). This relocation causes a drastic rearrangement of the apical loop, adopting an inactive conformation (**Fig. 4b**). This inactive conformation is stabilized by the hydrogen bonds between the residues in the apical loop (Gly251, Gly252, and Arg256) and rRNA nucleotides (**Fig. 4c**). In addition to the conformational change of the apical loop, we observed a notable shift of the entire domain III of RF2, approximately 3 Å toward the L1 stalk, in comparison with its position in the canonical termination complex (**Fig. 4a and 4b**). Residues within the shifted region (Arg245, Glu258, Arg262, Gln280, and His281) further stabilize this conformation through hydrogen-bonding interactions with rRNA nucleotides (**Fig. 4c, 4d, and** 4e). Since there are several hydrogen-bonding interactions between the domain III and rRNA that are absent in the canonical termination complex, the rearranged and shifted RF2 conformation observed in our structure could be functionally important for general termination regulation.

**Figure 4.**
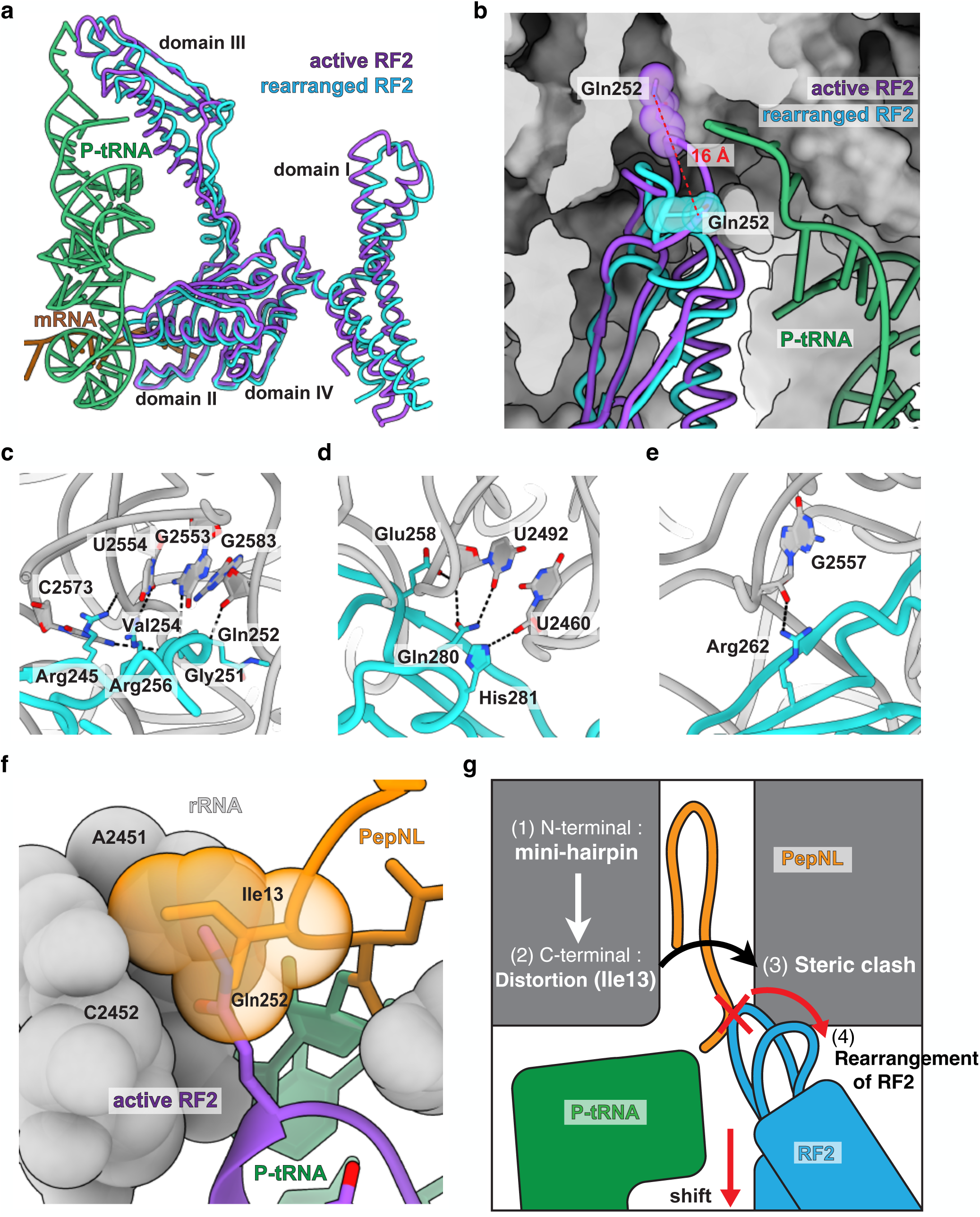
PepNL blocks RF2-mediated translation termination. **a.** The wild-type (UGA) or mutated (UAG) *pepNL* mRNA was translated by PURE*frex* in the presence or absence of tryptophan (Trp), and the PepNL-arrested ribosome was visualized by toeprint analysis. **b.** The wild-type (UGA) or mutated (UAG) *pepNL* mRNA was translated by PURE*frex* in the absence or presence of tryptophan. The ^35^S-methionine-labeled translation products were separated by neutral pH SDS-PAGE with optional RNase A (RN) pretreatment. Release factors (RFs) are omitted where indicated (lanes 11 and 12). The *pepNL* D14 nonstop mRNA (D14-NS) was also analyzed to serve as the positional marker (lanes 9 and 10). The PepNL (14 aa) or PepNL (read-through: RT) peptidyl-tRNA and PepNL (14 aa) or PepNL (RT) peptide were schematically indicated. An asterisk denotes the fMet-tRNA. **c.** The *pepNL*-stop (UGA/UAA/UAG)-*lacZ* mRNA was expressed in *E. coli* cells, and the frequency of stop codon read-through was calculated as described in the Materials and Methods. **d.** The wild-type *pepNL* (UGA) mRNA was translated by PURE*frex* in the absence of tryptophan and release factors (RF). Then 25 µM of tryptophan was added and further incubated for the indicated duration. **e.** The schematic illustration of tryptophan-dependent release of the PepNL-arrested ribosome. RF2 inefficiently terminates the translation of *pepNL* due to the steric clash shown in Fig. 4. However, in the presence of sufficient tryptophan, the Trp-tRNA competitively decodes the UGA of *pepNL*, leading to the stop codon read-through and resolution of PepNL-arrested ribosome.

### The PepNL nascent peptide inhibits RF2 activity through a steric clash with Ile13

Further structural comparison showed that the side chain of Ile13 in the distorted C-terminus of PepNL nascent peptide is accommodated in the pocket formed by A2451, C2452, and U2506 (**Fig. 4f**). This pocket is crucial for translation termination, as the ^250th^GGQ^252nd^ motif of RF2 enters this pocket to cleave the ester bond (**5–8, 57**). Consequently, the distortion of Ile13 shown in **Fig. 3i** leads to a steric clash with the Gln252 of RF2, hampering the accommodation of the GGQ motif in the A2451/C2452/U2506-forming pocket (**Fig. 4f**). This steric clash between Ile13 of PepNL and Gln252 of RF2 would induce a drastic rearrangement of the apical loop and subsequent rearrangement of the entire domain III of RF2. These considerations are supported by the finding that the Ile13 mutation abolished the arrest activity of PepNL (**Fig. 3j, Supplementary Fig. 3c**).

Taken together, our cryo-EM reconstruction revealed the molecular details of how PepNL nascent peptide inhibits the translation termination activity of RF2 (**Fig. 4g**). The translated nascent PepNL peptide forms a compact hairpin structure, with its N-terminus directed back to the PTC, and stacks inside the exit tunnel. This stacking distorts the C-terminus of the PepNL nascent peptide, leading to the steric clash between Ile13 of PepNL and Gln252 of RF2. Consequently, this steric clash induces a drastic conformational change in RF2, shifting it into an inactive conformation.

### Read-through of the stop codon acts as an arrest-release mechanism for PepNL

Despite our success in determining the structure of the PepNL-arrested ribosome, we initially faced difficulties in preparing the stalled ribosome within the complete PURE*frex* system (**Fig. 5a, lanes 1 and 2**). However, our experiments with the *pepNL* mutant, harboring a UAG stop codon instead of UGA, revealed robust ribosome stalling both *in vitro* and *in vivo* (**Fig. 5a, lanes 5 and 6, Supplementary Fig. 6a and 6b**). Furthermore, the *pepNL* mutants carrying altered stop codon exhibited increased cytotoxicity (**Supplementary Fig. 6c**). These results indicate that the UGA stop codon of *pepNL* plays a role in attenuating ribosome stalling. To investigate this, we conducted experiments where tryptophan, known to induce read-through at the UGA codon (**58–60**), was excluded from the *in vitro* translation. Remarkably, even with the wild-type *pepNL* (UGA), robust ribosome stalling was observed (**Fig. 5a, lanes 3 and 4**). Electrophoretic analysis of the polypeptide products revealed more significant accumulation of peptidyl-tRNA species in the absence of tryptophan (**Fig. 5b, lanes 1, and 5**). In addition, we observed another peptidyl-tRNA migrating more slowly than the full-length PepNL (14 aa)-tRNA, in the presence of tryptophan and the absence of release factors (**Fig. 5b, lanes 11 and 12**). These indicate that ribosomes read-through the UGA stop codon in the presence of tryptophan and stall at downstream stop codons (**Supplementary Fig. 6a**). Consistent with this assumption, our *in vivo* reporter assays demonstrated that the UGA codon, but not UAA or UAG, exclusively induced the read-through of the *pepNL* stop codon (**Fig. 5c**). Furthermore, tryptophan effectively induced the stop codon read-through even after the ribosome was arrested by PepNL (**Fig. 5d**), indicating that stalled ribosome is “released” by Trp-tRNA-mediated read-through (**Fig. 5e**).

**Figure 5.**
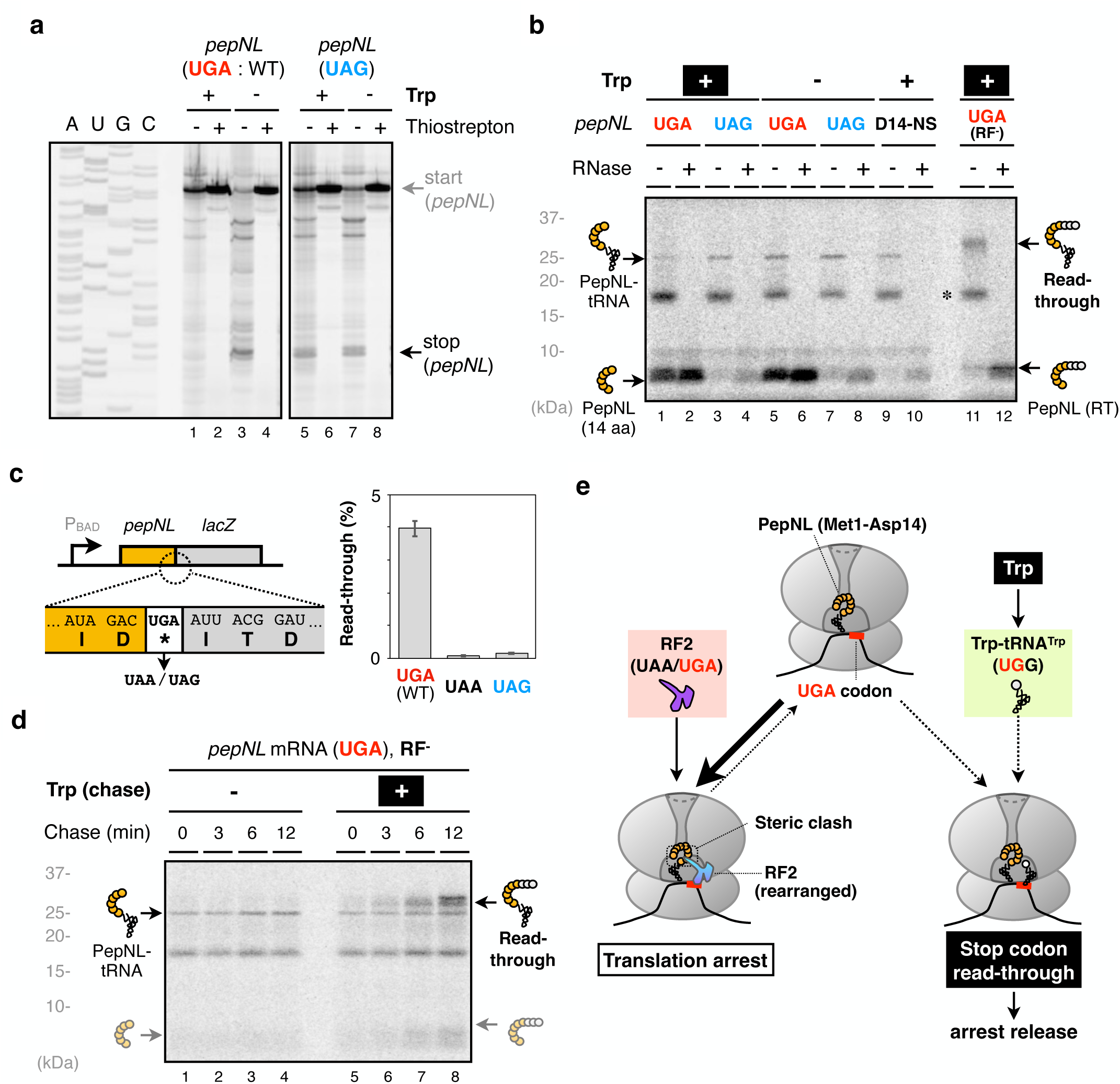
Read-through of stop codon acts as an arrest-release mechanism for PepNL. **a**, Overview of the RF2 structure within the PepNL-arrested ribosome (cyan) compared with the active RF2 (purple, PDB: 6C5L). The P-tRNA^Asp^ (green) and *pepNL* mRNA (brown) are also shown. **b**, Close-up view of the apical loop conformations of the rearranged and active RF2 states. Gln252s of the GGQ motif are highlighted as stick with sphere models. Ribosome is shown as a surface model. **c**, **d**, **e**, Hydrogen bond interactions between the apical loop of the rearranged RF2 and 23S rRNA. **f**, Superposition of the active RF2 (purple) indicates a steric clash of its Gln252 with Ile13 of PepNL. Distorted Ile13 of the PepNL peptide occupies the pocket formed by A2451, C2452, and U2506 (not shown) of 23S rRNA, blocking the proper accommodation of the RF2 apical loop. The rRNA nucleotides and Ile13 are shown as sphere models. **g**, Schematic representation of the inhibition mechanism of RF2 by the PepNL peptide.

## Discussion

Our study demonstrated that small ORFs *pepNL* and *nanCL* in *E. coli* encode ribosome arrest peptides (RAP) (**Fig. 1 and 2**). Furthermore, we successfully determined the cryo-EM structure of the ribosome arrested by PepNL (**Fig. 3 and 4**). Our structure revealed that the nascent PepNL peptide adopts a distinctive “mini hairpin” conformation near the entrance of the exit tunnel, causing the distortion of its C-terminal segment, including Ile13 residue. This distorted structure prevents the proper accommodation of the GGQ motif of RF2, inhibiting peptide release. Importantly, PepNL-dependent stalling does not require an “arrest inducer”, instead the stalling is resolved via stop codon read-through induced by tryptophan (**Fig. 5**).

The PepNL nascent peptide within the tunnel adopts a unique fold to enable a novel arresting mechanism by employing only 14 amino acid residues. Generally, the N-terminus of the nascent peptide elongates toward the exit of the tunnel (**9, 12, 56**). In contrast, in our structure, the N-terminus of PepNL is oriented in the opposite direction, toward the tunnel entrance. While it has been reported that nascent peptides of VemP and XBP1u adopt S-shaped conformation with partially folding back in the exit tunnel (**31, 32**), U-shaped conformation with overall reversal of the nascent peptide as in the present PepNL structure has not yet been reported. Moreover, the translation inhibition mechanism by this “mini hairpin” structure might raise a new concept at the early stage of protein synthesis. Previous studies have reported that N-terminal amino acid sequences immediately adjacent to the initiation codon can affect translation efficiency (**61, 62**). Stochastic formation of the mini hairpin-like structure could potentially inhibit the expression of proteins, which is not limited to PepNL.

Previous cryo-EM studies have elucidated the details of how arrest peptides prevent translation termination (**21, 22, 33**). In the case of TnaC, U2585, A2602 of 23S rRNA and Arg23 (Phe23) of TnaC nascent peptide cooperatively inhibit proper accommodation of RF2 (**22**). In SpeFL, the distortion of U2585 by Asn32 of SpeFL nascent peptide causes a steric clash with RF1, thereby inhibiting peptide release (**33**). In the present structure, the steric clash between Ile13 of PepNL nascent peptide and the GGQ motif of RF2 inhibits peptide release. These three RAPs commonly prevent the proper accommodation of the GGQ motif in distinct manners. Moreover, our study once again emphasizes the significance of the second-to-last amino acid residue (Ile13 in PepNL) for peptide release inhibition (**22, 63**). Earlier studies have pointed out that the efficiency of RF-mediated termination is affected by the amino acid sequence of the nascent peptide (**64–66**). Notably, Isaksson and colleagues pointed out the importance of the penultimate C-terminal residue in the canonical translation termination (**63**). The inhibition mechanisms revealed by recent structural analyses may contribute to a deeper understanding of the universal translation cycle. Our successful observation of the entire structure of distorted RF2 could offer several insights. We observed that the Ile13 of PepNL excluded the GGQ motif, causing a drastic rearrangement of the apical loop within domain III of RF2. In addition, this noncanonical structure of RF2 is stabilized by additional hydrogen bonds. Therefore, it’s plausible to consider the GGQ motif as a sensor to fine-tune translation termination efficiency in response to the structural state within the PTC. When there’s a mismatch between the GGQ motif and the PTC structure, it could result in the stabilization of the non-canonical RF2 conformation via additional interactions, potentially delaying the translation termination.

In contrast to sensory arrest peptides such as TnaC and SpeFL, the folding of PepNL nascent peptide within the tunnel is independent of the metabolite binding. Instead, PepNL utilizes the stop codon read-through as an “arrest releaser”. This framework makes PepNL a metabolite-sensing arrest peptide in a distinct way from the previously studied RAPs. Subsequently, the translation of *pepNL* then regulates the expression of the downstream PepN, depending on the availability of tryptophan. PepN, an endopeptidase, degrades not only oligopeptides but also full-length polypeptides with a broad substrate specificity (**67**). In nutrient-limited conditions, PepN has a crucial role in adaptation through the reproduction of free amino acids (**68**). These suggest that the correlation between PepN activity and the concentration of intercellular amino acids holds significance for environmental adaptation.

Our approaches identified two novel RAPs PepNL and NanCL that function in *E. coli* living cells. However, it should be noted that our evaluations for the RAP activity of each sORF are limited to the standard laboratory conditions (LB medium, 37 °C, aerobic condition). In fact, SpeFL, which responds to ornithine, did not exhibit a significant result under the conditions we tested, indicating the limitations of our approaches. On the other hand, our analysis did not exclude the possibility that sORFs other than *pepNL* and *nanCL* encode sensory arrest peptides. In our study, the translation of 19 out of 39 sORFs significantly influenced the expression of downstream genes, implying that they harbor translation-coupled functions (**Supplementary Fig. 1e**). Further analyses of these sORFs would expand our understanding of the translation regulations.

To explore the unidentified “arrest inducer” for each RAP candidate, monitoring the expression of Cold Shock Proteins (CSPs) could serve as a valuable indicator. While the stalling of translation elongation commonly triggers the expression of CSPs, the precise mechanism has not yet been elucidated (**43**). This enigmatic phenomenon should be studied in the future, however, if similar responses occur across species (**69, 70**), it could be a convenient method for evaluating RAP activity. In addition, this study also demonstrated the effectiveness of LC-MS/MS measurement of nascent peptidyl-tRNA species to identify RAPs. Our approaches to identify PepNL and NanCL, as well as the distinct molecular mechanism of translation stalling and regulation, have provided valuable insights into deciphering the hidden genetic codes within polypeptide sequences.

## Materials and methods

### *E. coli* strains, plasmids, and primers

*E. coli* strain BW25113 {Δ(*araD-araB*)567, Δ*lacZ4787*(::*rrnB*-3), λ-, *rph*-1, Δ(*rhaD*-*rhaB*)568, *hsdR514*} was used as the experimental standard strain. Plasmids used in this study are listed in **Supplementary Table 2.** Plasmids were constructed using standard cloning procedures, including Gibson assembly and QuickChange PCR. The sequence files of the plasmids constructed in this study are available in the Mendeley repository (doi: 10.17632/2bkz2xnnn5.1).

### Spot assay

*E. coli* cells harboring pCA24N with each sORF were grown overnight at 37 °C in LB medium supplemented with 20 µg/ml chloramphenicol. On the next day, they were serially diluted with fresh LB medium (10^-1^, 10^-2^, 10^-3^, 10^-4^, 10^-5^, and 10^-6^) and spotted onto LB agar plates containing 20 µg/ml chloramphenicol, with or without 1 mM IPTG (isopropyl β-D-thiogalactopyranoside, Nacalai tesque). The plates were incubated at 37 °C for 1 overnight, and the growth of colonies was recorded.

### β-galactosidase assay

*E. coli* cells harboring the *lacZ* reporter plasmid were grown overnight at 37 °C in LB medium supplemented with 100 µg/ml ampicillin. On the next day, the cultures were inoculated into fresh LB medium containing 2 x 10^-3^% arabinose and 100 µg/ml ampicillin, and were grown at 37 °C until the A_660_ reached 0.5. Afterward, 20 µl portions were subjected to a β-galactosidase assay to caluculate the Miller unit (m.u.) as described (**71**). The average of three independent experiments is presented with a standard error (SE) value.

### uORF-dependent regulation score

The uORF-dependent regulation score was calculated in the following formula.

uORF-dependent regulation score

= {m.u. (Plasmid-B)} / [{m.u. (Plasmid-B)} + m.u. (Plasmid-A)}]

#### Plasmid-A

A plasmid carrying P_BAD_ promoter, 5’ region of the downstream main ORF (mORF) relative to the small ORF, spanning from the transcription start site to the initiation codon of the mORF, and *lacZ* reporter which is translationally fused with the mORF.

#### Plasmid-B

A derivative of plasmid-A with a mutation (ATG to ACG) to disrupt the translation initiation codon (AUG) or with insertion of a stop codon to prematurely terminate the translation of the small ORF. Detailed information on these mutations can be found in **Supplementary Table 2**, and sequence files are available in the Mendeley repository (doi: 10.17632/2bkz2xnnn5.1).

### The frequency of stop codon read-through

*E. coli* cells expressing the *pepNL*-stop-*lacZ* or its derivative lacking the stop codon of *pepNL* reporter were grown and subjected to a β-galactosidase assay. Then the frequency of stop codon read-through was calculated in the following formula.

### Stop codon read-through (%)

= m.u. (*pepNL*-stop-*lacZ*) / m.u. (*pepNL*Δstop-*lacZ*)

### *In vitro* translation and product analysis

The coupled transcription-translation reaction was performed using PURE*frex* v1.0 (GeneFrontier) in the presence of ^35^S-methionine {EasyTag L-[^35^S]-Methionine (PerkinElmer)} at 37 °C, as described previously (**38**). DNA templates were prepared by PCR, as summarized in **Supplementary Table 3**. Tryptophan or release factors (RF: RF1, RF2, RF3, and RRF) were excluded from the *in vitro* translation mixture if indicated. The reaction was stopped by the addition of ice-cooled 5% TCA, washed by ice-cooled acetone, dissolved in SDS sample buffer (125 mM Tris-HCl, pH 6.8, 2% SDS, 10% glycerol, 50 mM DTT) that had been treated with RNAsecure (Ambion). Finally, the sample was divided into two portions, one of which was incubated with 50 µg /ml of RNase A (Promega) at 37 °C for 60 min, and separated by a WIDE RANGE Gel SDS-PAGE system (Nakalai Tesque). Radioactive bands were developed by using an imaging plate and Amersham™ Typhoon™ scanner RGB system (GE Healthcare).

### Toeprint analysis

Toeprint analysis was performed as described previously (**38, 72**). *In vitro* translation reaction sample was mixed with an equal volume of reverse transcription mixture [50 mM HEPES-KOH pH 7.6, 100 mM potassium glutamate, 2 mM spermidine, 13 mM magnesium acetate, 1 mM DTT, 2 µM fluorescently labeled oligonucleotide (pe-lacZ-N-rv with Alexa 647 at 5’-terminus), 40 µM of each of dNTPs, 10 unit/µl ReverTra Ace (Toyobo)] and incubated at 37 °C for 10 min. The reverse transcription products were purified by NucleoSpin Gel and PCR clean-up kit equilibrated with NTC buffer (Macherey-Nagel). Dideoxy DNA sequencing samples were prepared using Thermo Sequenase DNA polymerase (Cytiva) and the same templates and primer (pe-lacZ-N-rv) as used for toeprint analysis. Samples were subjected to 8 % polyacrylamide-7 M urea-TBE gel electrophoresis. Fluorescent images were visualized and analyzed by Amersham™ Typhoon™ scanner RGB system (GE healthcare) using a 635 nm excitation laser and LPR emission filter. The final 100 µg/ml of thiostrepton was added beforehand if indicated. Tryptophan or release factors (RFs: RF1, RF2, RF3, and RRF) were excluded from the *in vitro* translation mixture if indicated.

### Northern blotting

*E. coli* cells were grown in LB medium until the A_660_ reached ∼0.5. The expression of *pepNL* was then induced by 1 mM IPTG for 10 min. After centrifuging and collecting the bacterial cells, total RNA was extracted using TriPure Isolation Reagent (Roche), according to the supplier’s instructions. RNA samples were dissolved in SDS sample buffer, separated by 11 % WIDE RANGE Gel SDS-PAGE, transferred onto BrightStar™-Plus Positively Charged Nylon Membrane (Invitrogen^TM^), and hybridized with a biotinylated oligonucleotide complementary to the tRNA^Asp^; CGGAACGGACGGGACTCGAACCCGCGACC. Hybridization experiments were performed using the ULTRAhyb™ Ultrasensitive Hybridization Buffer (Invitrogen^TM^) and Chemiluminescent Nucleic Acid Detection Module (Thermo Scientific) according to the manufacturer’s instructions. Images were visualized and analyzed by LAS4000 LuminoImager (GE Healthcare).

### Preparation of the lysate for proteomic analysis

Samples for proteomic analysis were prepared as described previously (**73**). *E. coli* cells harboring pCA24N with each sORF were grown in LB medium supplemented with 20 µg/ml of chloramphenicol until the A_660_ reached ∼0.2. Subsequently, 1 mM of IPTG was added to induce the expression of small ORFs, and cells were further incubated for 1h with shaking at 37 °C. The cells were then harvested and resuspended with PBS buffer (137 mM NaCl, 8.1 mM Na_2_HPO_4_, 2.68 mM KCl, 1.47 mM KH_2_PO_4_, pH 7.4). The suspension was mixed with an equal volume of 10% of TCA. After standing on ice for at least 10 min, the samples were centrifuged, and the supernatant was removed by aspiration. Precipitates were washed twice with acetone, by vigorous mixing. Proteins were dissolved in PTS buffer (12 mM sodium deoxycholate, 12 mM sodium lauryl sulfate, 100 mM Tris-HCl, pH 9.0), and 50 µg of total protein at a concentration of 1 µg/µl was processed to reduction by dithiothreitol (DTT), alkylation with iodoacetamide, and limited digestion by Trypsin/Lys-C Mix (Promega). The samples were then extracted by ethyl acetate, evaporated, dissolved in MS buffer-A (0.1 % TFA and 2 % acetonitrile), desalted by StageTip composed of an SDB-XC Empore disk (3M, U.S.A.), eluted with MS buffer-B (0.1 % TFA and 80 % acetonitrile), again evaporated, and resolved in MS buffer-A. Finally, the samples were subjected to the LC-MS/MS measurement.

### LC-MS/MS analysis for a comparative quantitative proteomic analysis

In a series of quantitative proteomic analyses, we used the Eksigent NanoLC 415 nanoflow HPLC system and the TripleTOF 4600 tandem-mass spectrometer (AB Sciex, U. S. A.) in DIA/SWATH acquisition mode (**74**). The detailed settings for the LC-MS/MS measurements are summarized in **Supplementary Table 4**. The measurement was conducted three times for each sample.

Data analysis was performed by the DIA-NN software (version 1.8.1, https://github. com/vdemichev/diann, downloaded on 1 November 2023) (**75**). The spectral library for DIA/SWATH analysis was obtained from the SWATH atlas (http://www.swathatlas.org/, accessed on 30 April 2021); the original data were acquired by Midha *et al*. (**76**). All downstream statistical analyses were performed using in-house R scripts (R.app for Mac, version 4.3.1). Only the proteins with intensities obtained in all three measurements in both samples were used to calculate fold changes. *P*-values for the volcano plots were calculated using Welch’s t-test with the Benjamini-Hochberg correction (using the “p.adjust” function in R.app). PCA and hierarchical clustering analyses were performed using the data from 1,205 proteins for which foldchange values were obtained under all 36 conditions.

### Preparation of the peptidyl-tRNA for LC-MS/MS analyses

*E. coli* cells harboring pCA24N with each sORF were grown in LB medium supplemented with 20 µg/ml of chloramphenicol until the A_660_ reached approximately 0.5. Subsequently, 1 mM of IPTG was added to induce the expression of small ORFs, and cells were further incubated for 30 min with shaking at 37 °C. Afterward, the cell culture was mixed with an equal volume of ice-cooled 10% TCA to precipitate the cellular components. The precipitate was washed twice with ice-cooled acetone, and dissolved in PTS buffer.

The peptidyl-tRNAs in the PTS buffer were isolated by using a High Pure miRNA Isolation Kit (Roche), as described previously (**54**). After adding *binding buffer* (provided by the manufacturer), the lysate was vortexed and mixed at 37 °C for 30 min with shaking. Then *binding enhancer* (provided by the manufacturer) was added and the lysate was centrifuged at 12,000 × g for 3 min at 4 °C to remove the precipitates. The supernatant was loaded into silica columns, and the following steps were performed according to the manufacturer’s instructions. The silica-bound RNA was eluted with PTS buffer. This purification was repeated twice.

The purified RNA sample was mixed with 0.1x volume of 2 M Tris base (pH= ∼11) and incubated at 80 °C for 20 min to hydrolyze the ester bond of the peptidyl-tRNA. After the addition of 0.9x volume of deionized water, the sample was reduced by DTT, alkylated with iodoacetamide, and digested with Lys-C (Fujifilm-Wako) or Lys-C / Trypsin mix (Promega). Then samples were extracted by ethyl acetate, evaporated, dissolved in MS buffer-A (0.1 % TFA and 2 % acetonitrile), desalted by StageTip composed of an SDB-XC Empore disk, eluted with MS buffer-B (0.1 % TFA and 80 % acetonitrile), again evaporated, and resolved in MS buffer-A.

### LC-MS/MS analysis for peptides derived from peptidyl-tRNAs

For the detection of the peptides derived from peptidyl-tRNAs, the Easy-nLC 1000 nanoflow HPLC system and the Q-Exactive tandem-mass spectrometer (Thermo Fisher Scientific, U. S. A.) were used in data-dependent acquisition (DDA) mode. The detailed settings for the LC-MS/MS measurements are summarized in **Supplementary Table 5**. The data analysis of the DDA measurements was performed with the Proteome Discoverer 2.4 software bundled with the SEQUEST HT search engine (Thermo Fisher Scientific, U. S. A.). To detect stalled peptides in translation, Trypsin (semi) or LysC (semi) setting was used for enzymatic digestion.

### Statistical analyses

Statistical analyses were conducted by using R (https://www.r-project.org) and Python (https://www.python.org).

### GWIPS-viz browser

GWIPS-viz browser (**42**) was utilized with the following settings.

Organisms : *Escherichia coli* K12

Elongating Ribosome (A-site): Global Aggregates.

mRNA-seq Reads: Global Aggregates.

### Grid preparation and data acquisition

For the cryo-EM analysis, 3 µL of the PURE system reaction solution containing stalled ribosomes was applied onto a glow-discharged holey-carbon grid coated with continuous 2 nm-thick carbon film (Quantifoil Au 300 mesh, R2/1 + 2 nm C) and incubated for 30 s in a controlled environment of 100% humidity at 4 ℃. The grids were blotted for 4 s, followed by plunge freezing in liquid ethane, using a Vitrobot Mark IV (FEI).

The datasets were collected using a Titan Krios G4 microscope (Thermo Fisher Scientific), running at 300 kV and equipped with a Gatan Quantum-LS Energy Filter (GIF). Gatan K3 Summit direct electron detector was used at a pixel size of 0.83 Å (magnification of ×105,000) with a dose of approximately 30 electrons per Å^2^ with 30 frames. The data were automatically acquired using the EPU software (Thermo Fisher Scientific), with a defocus range of -0.8 to -1.6 µm, and 15,342 movies were obtained. Detailed parameters are listed in **Supplementary Table 6**.

### Image processing

The data were processed by the cryoSPARC v4.4.0 software platform (**77**). The dose-fractionated movies were subjected to beam-induced motion correction and dose weighting using patch motion correction, and the contrast transfer function (CTF) parameters were estimated using patch-based CTF estimation. From the 15,342 preprocessed micrographs, 2,521,444 particles were automatically picked by the template picker, using the references created by the blob picker and several rounds of 2D classification. The particles were subjected to several rounds of reference-free 2D classifications to create particle sets. The curated particles were aligned by non-uniform refinement (**78**) using the *E. coli* ribosome map (EMD-22586) as the reference, followed by unsupervised 3D classification and a successive 2D classification to eliminate the ratcheted ribosomes and junk particles. The remaining particles were further classified by no-align focused 3D classification using the mask covering P-site, A-site, and the binding sites of EF-Tu:tRNA and RF2, resulting in four classes with RF2 bound, A-tRNA bound, EF-Tu:tRNA bound, and empty ribosomes, respectively. The particles of the RF2-bound ribosome (71,980 particles) were subjected to CTF refinement (beam-tilt, trefoil, fourth-order aberrations, and per-particle defocus), reference-based motion correction, followed by the final non-uniform refinement. The global resolution of the final map is 2.9 Å based on the gold-standard, applying the 0.143 criterion on the Fourier shell correlation (FSC) between the reconstructed half maps (**79**).

### Model building and refinement

The reported high-resolution model of *E. coli* ribosome (PDB 7k00, **reference-80**), including bound ligands and metal ions, was used as the starting model, while that of RF2 was from 6OUO. The models were docked into our map by rigid body fit, followed by jiggle fit (ref) and manual revision in Coot (**81**). The pepNL peptide was built manually based on the density. Water molecules were automatically picked by *Coot*, followed by manual revision. Metal ions were revised and added based on the coordination pattern and the density, which is stronger in the map without sharpening due to their positive charge. Geometrical restraints of modified residues and ligands were calculated by Grade Web Server (http://grade.globalphasing.org). The final model was subjected to the refinement of energy minimization and atomic displacement parameters (ADP) estimation by Phenix.real_space_refine v1.20 (**82**) with Ramachandran and rotamer restraints, against the map with *B*-factor sharpening and local-resolution filtering by cryoSPARC. Metal-coordination restraints were generated by ReadySet implemented in PHENIX and non-canonical covalent bond restraints for modified residues were manually prepared. The refined model was validated with MolProbity (**83**) and EMRinger (**84**) in the PHENIX package. Model refinement statistics are listed in **Supplementary Table 6**. UCSF ChimeraX 1.6 (**85**) was used to make the figures.

## Data Availability

Raw data files are available in the Mendeley Data repository (doi: 10.17632/2bkz2xnnn5.1). The mass spectrometry data have been deposited in the jPOST repository (<u>JPST003118 / PXD052430)</u> (**86**).

## Acknowledgement

We thank Takashi Kanamori for providing customized PURE*frex* v1.0. Additionally, we extend our thanks to the Center for Integrative Biosciences and the Biomaterials Analysis Division, the Open Facility Center at the Tokyo Institute of Technology for DNA sequencing, and the Cell Biology Center Research Core Facility at Tokyo Tech for the TripleTOF 4600 mass spectrometry measurements.

## Funding and additional information

This work was supported by MEXT Grants-in-Aid for Scientific Research (Grant Numbers JP20H05925 to H.T.), Research Support Project for Life Science and Drug Discovery (Basis for Supporting Innovative Drug Discovery and Life Science Research (BINDS)) from AMED under Grant Number JP22ama121012 (support number 4857 to H.T.) and JP23ama121002 (support number 3273 to O.N.), Japan Society for the Promotion of Science (JSPS) KAKENHI (Grant Number 22H02553 to Y.I., 23H02410 to Y.C.), and Core Research for Evolutional Science and Technology (CREST) of the Japan Science and Technology Agency (JST) (Grant Number JPMJCR20E2 to O.N.), and the grants from the Ohsumi Frontier Science Foundation, the Japan Foundation for Applied Enzymology, the Takeda Science Foundation, the Yamada Science Foundation and the Senri Life Science Foundation to Y.C.

## Author Contributions

A.K., Y.C., S.K., and Y.K. performed genetic and biochemical experiments. T.N. and A.Y. performed mass-spectrometry measurements and data analyses. Y.A. collected the cryo-EM data and built the model. Y.A. and Y.I. analyzed the structure. Y.C., Y.A., A.K., T.N., Y.I., O.N., and H.T. wrote the manuscript. Y.C., Y.I., H.T., and O.N. supervised the research.

