## Supplementary Information for "A newly-identified mini-hairpin shaped nascent peptide blocks translation termination by a novel mechanism"

**Supplementary Figures 1-6**

**Supplementary Dataset 1 and 2** (provided in separate files)

**Supplementary Tables S1-S4**

**Supplementary Table 1**: Peptide fragments identified by the LC-MS/MS analysis of peptidyl-tRNA (provided in a separate file).
**Supplementary Table 2**: Plasmids used in this study. (provided in a separate file)
**Supplementary Table 3**: Oligonucleotides used in this study. (provided in a separate file)
**Supplementary Table 4**: Parameters for SWATH acquisition by TripleTOF 4600.
**Supplementary Table 5**: Parameters for DDA measurement by Q-Exactive.
**Supplementary Table 6**: Data collection, processing, model refinement and validation statistics.

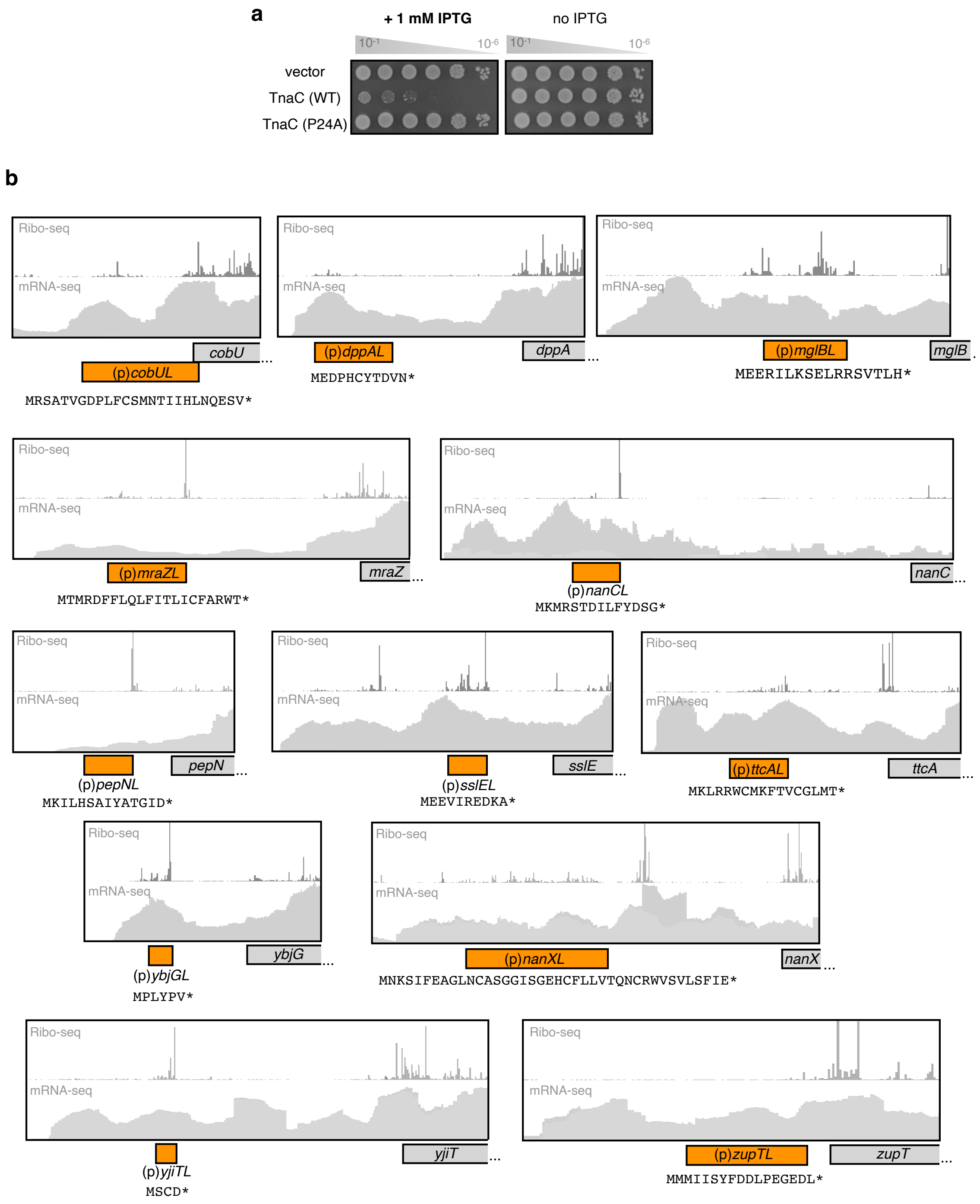
a
+ 1 mM IPTG
no IPTG
10-1
10-6
vector
TnaC (WT)
TnaC (P24A)
b
Ribo-seq
mRNA-seq
cobU
(p)cobUL
MRSATVGDPLFCSMNTIIHLNQESV*
(p)dppAL
dppA
MEDPHCYTDVN*
(p)mglBL
mglB
MEERILKSELRRSVTLH*
(p)mraZL
mraZ
MTMRDFFLQLFITLICFARWT*
(p)nanCL
nanC
MKMRSTDILFYDSG*
(p)pepNL
pepN
MKILHSAIYATGID*
(p)sslEL
sslE
MEEVIREDKA*
(p)ttcAL
ttcA
MKLRRWCMKFTVCGLMT*
(p)ybjGL
ybjG
MPLYPV*
(p)nanXL
nanX
MNKSIFEAGLNCASGGISGEHCFLLVTQNCRWVSVLSFIE*
(p)yjiTL
yjiT
MSCD*
(p)zupTL
zupT
MMMIISYFDDLPEGEDL*

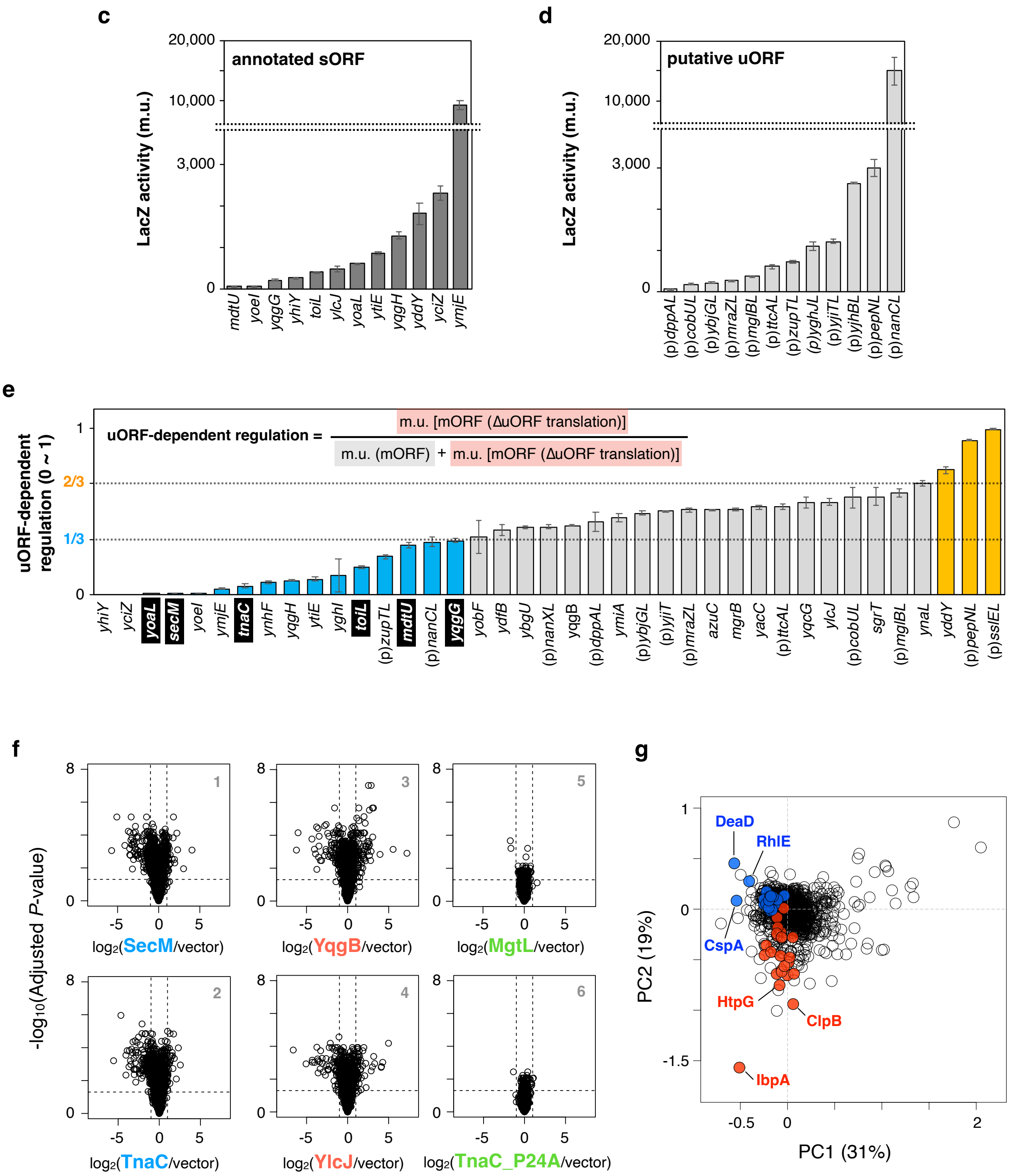

c
annotated sORF
LacZ activity (m.u.)
20,000
10,000
3,000
0
mdtU yoeI yqgG yhiY toiL ylcJ yoaL ytiE yqgH yddY yciZ ymjE
d
putative uORF
LacZ activity (m.u.)
20,000
10,000
3,000
0
(p)dppAL (p)cobUL (p)ybjGL (p)mraZL (p)mglBL (p)ttcAL (p)zupTL (p)yghJL (p)yjiTL (p)yjhBL (p)pepNL (p)nanCL
e
uORF-dependent regulation (0 ~ 1)
uORF-dependent regulation = m.u. [mORF (ΔuORF translation)] / m.u. (mORF) + m.u. [mORF (ΔuORF translation)]
1
2/3
1/3
0
yhiY yciZ yoaL secM yoeI ymjE tnaC ynhF yqgH ytiE yghI toiL (p)zupTL mdtU (p)nanCL yqgG yobF ydfB ybgU (p)nanXL yqgB (p)dppAL ymiA (p)ybjGL (p)yjiT (p)mraZL azuC mgrB yacC (p)ttcAL yqcG ylcJ (p)cobUL sgrT (p)mglBL ynaL yddY (p)pepNL (p)sslEL
f
-log10(Adjusted P-value)
8
0
-5 0 5
1 log2(SecM/vector)
2 log2(TnaC/vector)
3 log2(YqgB/vector)
4 log2(YlcJ/vector)
5 log2(MgtL/vector)
6 log2(TnaC_P24A/vector)
g
PC2 (19%)
PC1 (31%)
1
0
-1.5
-0.5
2
DeaD
RhlE
CspA
HtpG
ClpB
IbpA

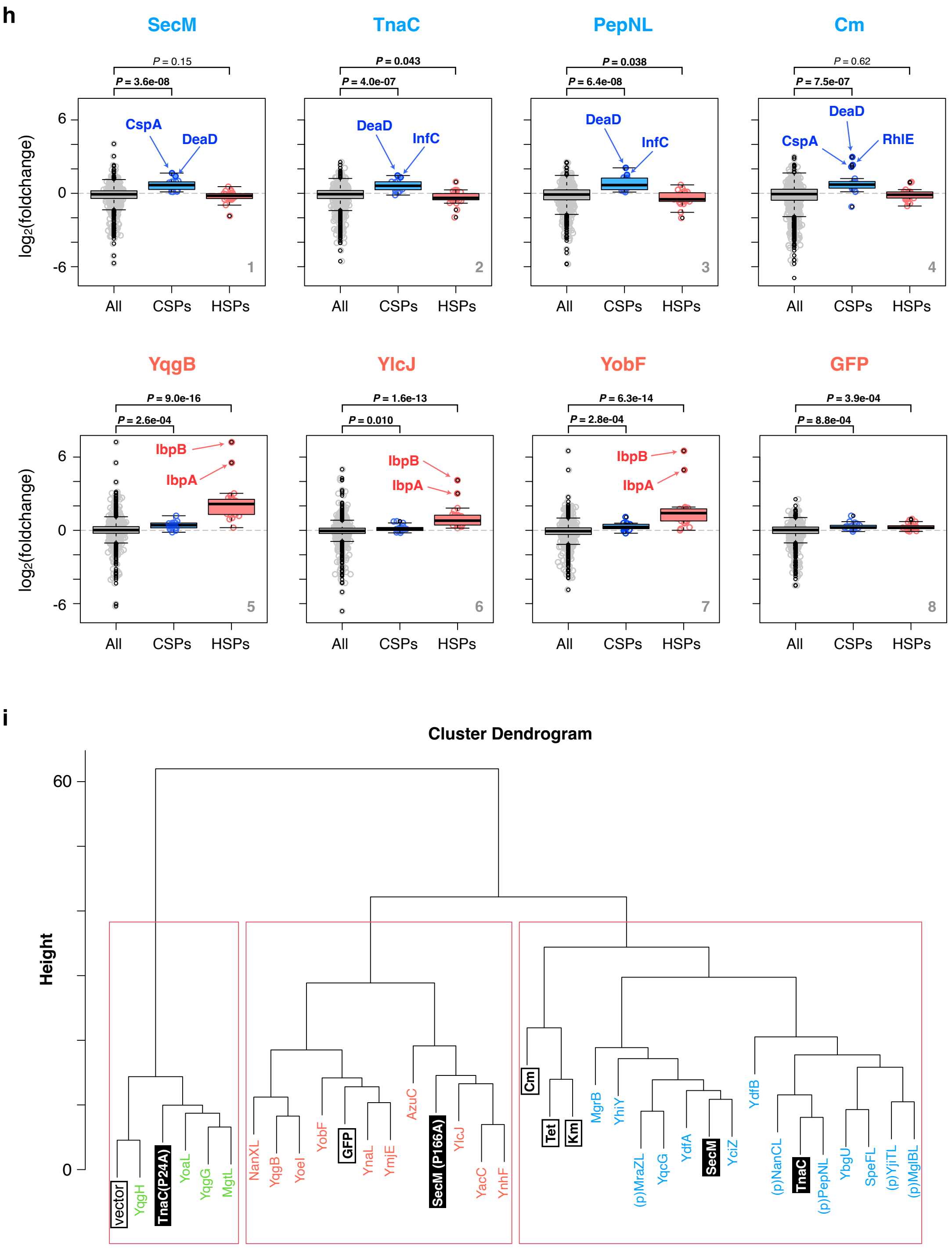


**Supplementary Figure 1.**

**a**. Serial dilution assay for evaluating the cytotoxicity when TnaC or TnaC (P24A) mutant was over-expressed in *E. coli* cells. A representative of three independent experiments is shown.

**b**. Putative small ORFs identified by using GWIPS-vis browser (**42**).

**c and d**. Translation efficiency of newly identified small ORFs. Each sORF, spanning from the transcription start site to its last coding codon, was cloned between the $P_{BAD}$ promoter and *lacZ* reporter. The expression of the LacZ reporter was quantified in Miller units (m.u.), and their mean values with standard errors (S.E.) were plotted. **C**: Previously annotated sORFs. **d**: putative sORFs identified in **Supplementary Fig. 1b**.

**e**. Regulatory function of sORFs. The 5' region of the downstream main ORF (mORF) relative to the small ORF, spanning from the transcription start site to the initiation codon of the mORF, was cloned between the $P_{BAD}$ promoter and *lacZ* reporter. The uORF-dependent regulation score was calculated as described in Materials and Methods. The mean values ±SE estimated from three independent biological replicates are shown. The mORFs whose expression is downregulated or upregulated in the absence of sORF translation are colored blue or orange, respectively.

**f**. Volcano plots of representative samples in the three populations. Colors (green, blue, and red) correspond to **Supplementary Fig. 1i**.

**g**. PCA loading plot of PC1 and PC2. Blue and red dots show major CSPs and HSPs, respectively.

**h.** Distributions of CSPs and HSPs in the representative samples in the two populations separated along with the PC2 axis. The numbers above the graph represent *P*-values by the Wilcoxon rank sum test against all evaluated data.

**i**. Dendrogram of a hierarchical clustering analysis of fold change values against a vector control. The number of clusters was set to three.

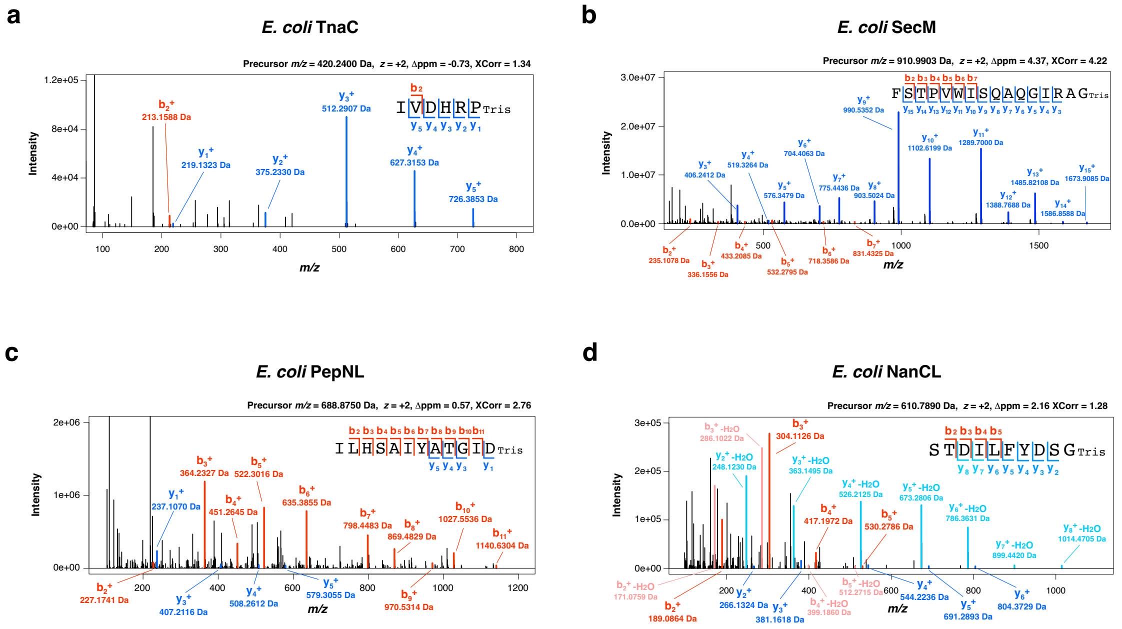


### Supplementary Figure 2.

**a-d.** MS/MS spectra of Tris-adducted peptides derived from peptidyl-tRNAs. The peaks of the b- and y- fragment ions are shown in red and blue, respectively. In **d**, the dehydrated y-ion fragments were also depicted since they appeared as major peaks.

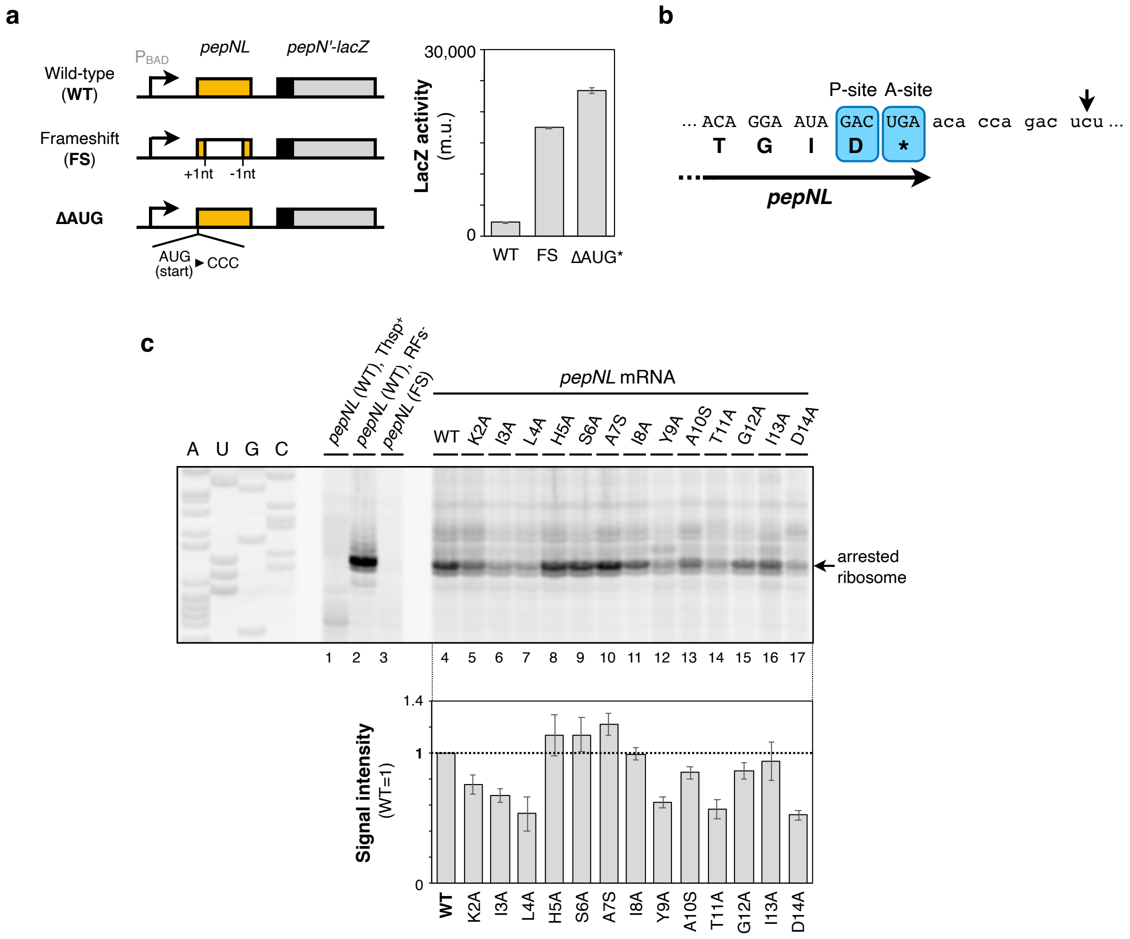


**Supplementary Figure 3.**

**a.** The *lacZ* reporter, fused with *pepN* and its 5' UTR containing the *pepNL* ORF, along with its derivatives carrying frameshift mutations or ATG to CCC initiation codon mutations, was expressed in *E. coli* cells. Subsequently, the expressed LacZ was quantified as Miller unit (m.u.), as described in the Materials and Methods. The mean values ±SE estimated from three independent biological replicates are shown.

**b.** Schematic illustration of the ribosome stalling site on the *pepNL* mRNA. The arrow indicates the point of reverse transcription interference, and the ribosomal occupancy is shown by the A-site and P-site codons.

**c.** The wild-type *pepNL* mRNA (lane 1, 2, and 4) or its varinats carrying frameshift mutation (lane 3) or substitution of the indicated amino acid (lanes 4-17) was translated by PURE*frex* in the absence of tryptophan, and the PepNL-arrested ribosome was visualized by toeprint analysis. Thiostrepton was pre-included (lane 1) or release factors ($RF^-$) was excluded from the reaction mixture (lane 2), if indicated.

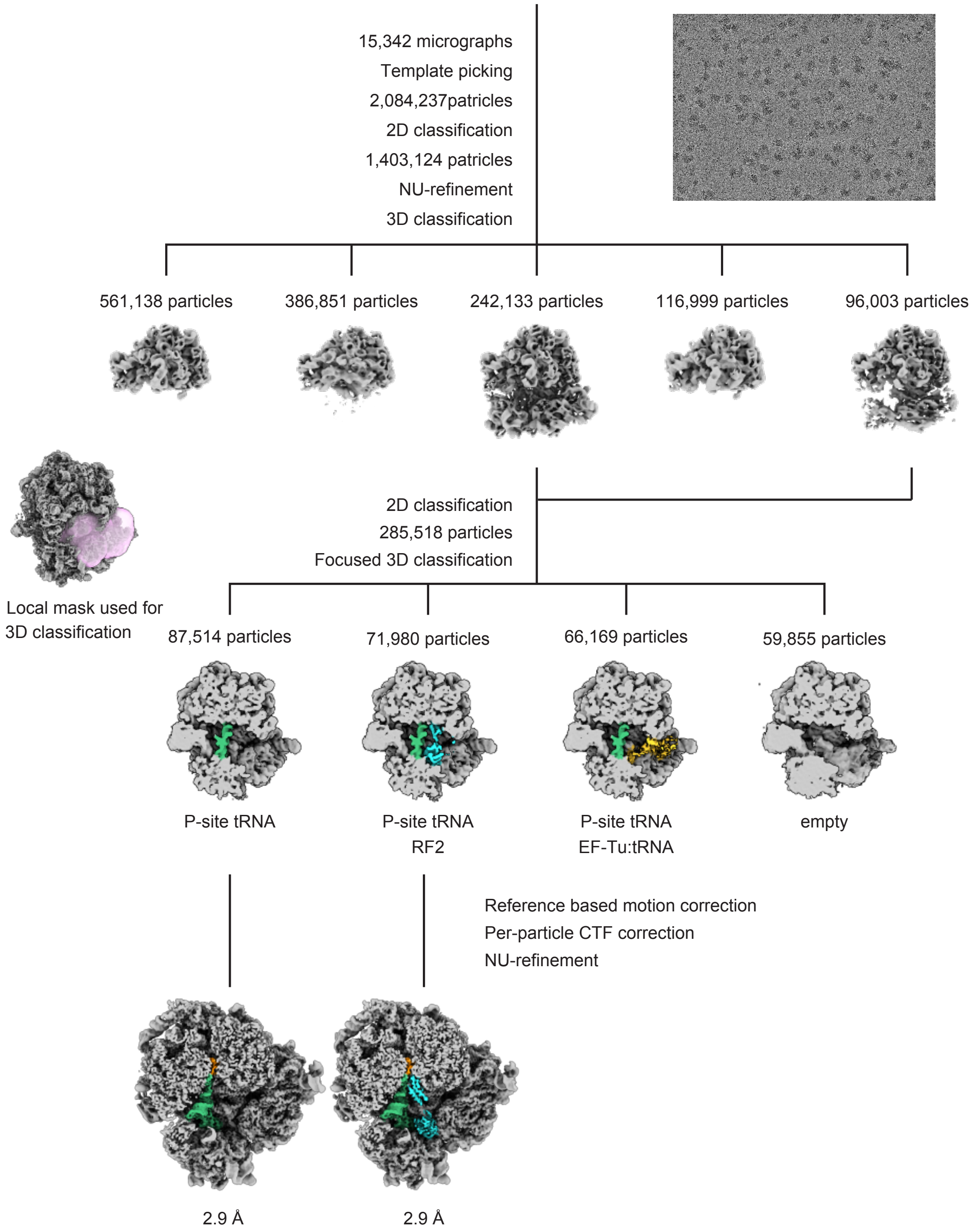


**Supplementary Figure 4.**

Single-particle cryo-EM image processing workflow for the final structure of the PepNL-70S ribosome complex in the presence or absence of RF2.

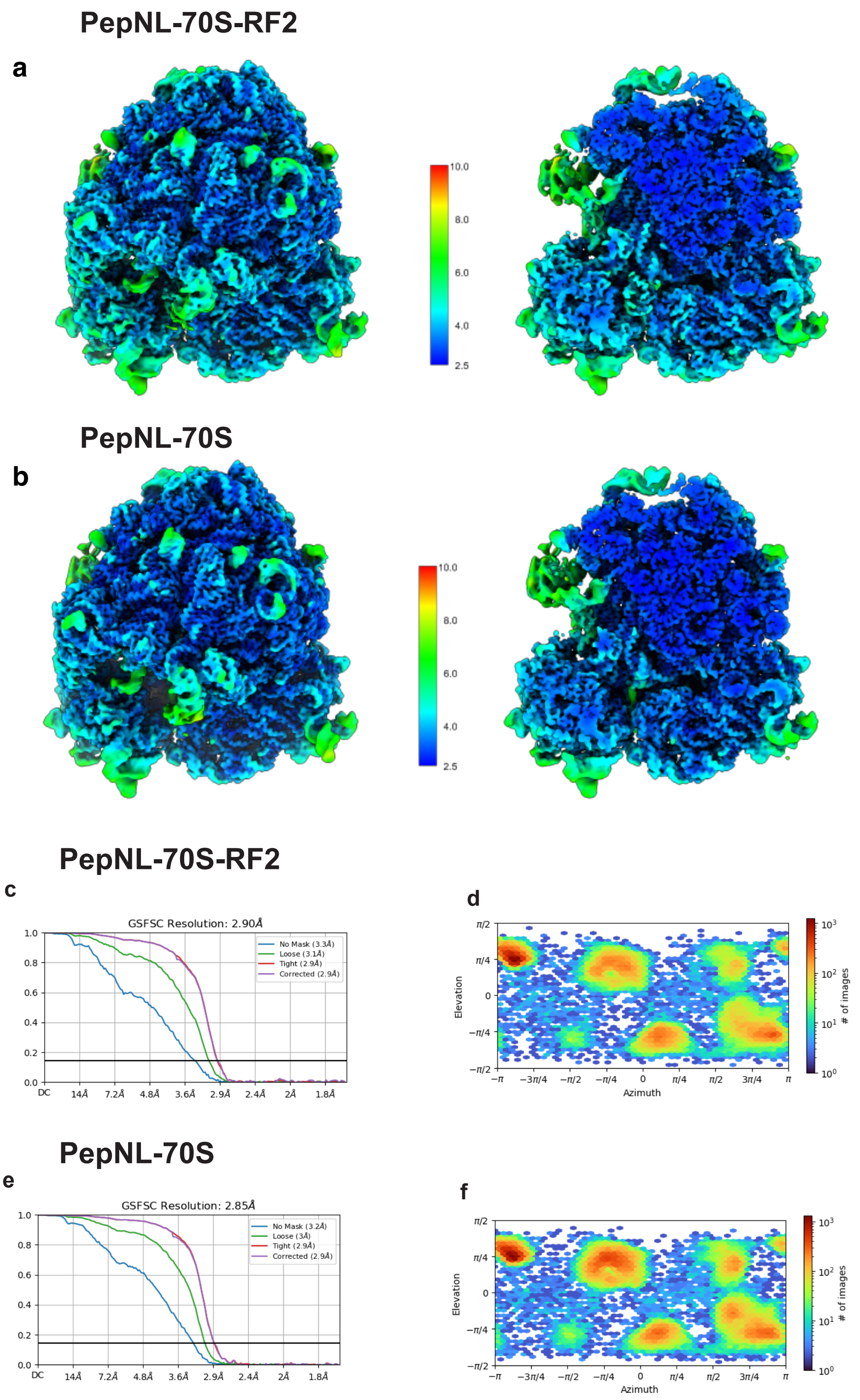


**Supplementary Figure 5.**

**a,b.** Local-resolution cryo-EM maps.

**c, d.** Fourier shell correlation (FSC) curves for the 3D reconstruction.

**e,f.** Direct distribution plot. (viewing distribution plot)

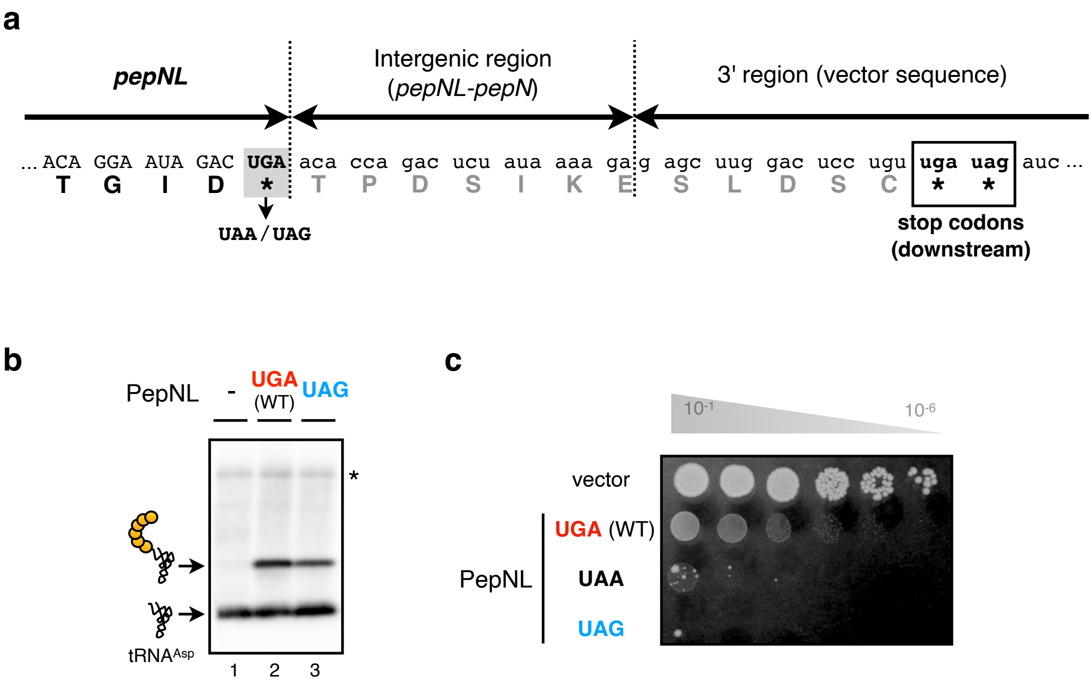


**Supplementary Figure 6.**

**a.** The nucleotide sequence of wild-type *pepNL* (WT: UGA) or its derivatives carrying altered stop codon (UAA or UAG). The downstream in-frame stop codons are also indicated.

**b.** The PepNL-tRNA accumulated in *E. coli* cells expressing the indicated *pepNL* variants was detected by northern blotting using anti-tRNA$^{Asp}$ probe.

**c.** Serial dilution assay to assess the cytotoxicity upon over-expression of wild-type PepNL (WT: UGA) or its derivatives carrying altered stop codon (UAA or UAG) in *E. coli* cells.

**Supplementary Table 4.** Parameters for SWATH acquisition by TripleTOF 4600.

| Parameter | | Value |
|---|---|---|
| nano HPLC | Trap column | 5.0 mm × 0.3 mm ODS column (L-column2, CERI, Japan) |
| | Separation column | 12.5 cm × 75 µm capillary column packed with 3 µm C18- silica particles (Nikkyo Technos, Japan) |
| | Flow rate | 300 nL/min |
| | Acetonitrile gradient (in the presence of 0.1 % formic acid) | 10 - 40%, 67 min |
| MS1 | Scan range | 350 - 1250 m/z |
| | Accumulation time | 100 msec |
| MS2 | Scan range (all window) | 350 - 1000 m/z |
| | Accumulation time | 60 msec |
| | Number of SWATH window | 40 |
| | m/z of SWATH window (variable window optimized for *E. coli* cell lysate) | 349.5 - 365.8<br>364.8 - 381.1<br>380.1 - 394.1<br>393.1 - 406.3<br>405.3 - 417.1<br>416.1 - 427.4<br>426.4 - 437.8<br>436.8 - 447.2<br>446.2 - 457.1<br>456.1 - 466.6<br>465.6 - 476.5<br>475.5 - 486.4<br>485.4 - 496.3<br>495.3 - 505.7<br>504.7 - 515.6<br>514.6 - 525.1<br>524.1 - 534.5<br>533.5 - 544.0<br>543.0 - 553.0<br>552.0 - 562.5<br>561.5 - 571.9<br>570.9 - 581.4<br>580.4 - 590.8<br>589.8 - 600.7<br>599.7 - 611.1<br>610.1 - 621.4<br>620.4 - 632.2<br>631.2 - 643.5 |

| | | |
|---|---|---|
| | | 642.5 - 655.2<br>654.2 - 668.2<br>667.2 - 681.7<br>680.7 - 697.0<br>696.0 - 714.1<br>713.1 - 732.6<br>731.6 - 754.6<br>753.6 - 780.7<br>779.7 - 812.7<br>811.7 - 851.4<br>850.4 - 906.7<br>905.7 - 999.9 |

**Supplementary Table 5.** Parameters for DDA measurement by Q-Exactive.

<table>
<tr><th colspan="2">Parameters</th><th>Values</th></tr>
<tr><td rowspan="4">nano HPLC</td><td>Trap column</td><td>2 cm × 75 µm capillary column<br>packed with 3 µm C18-silica particles<br>(Thermo Fisher Scientific, U. S. A.)</td></tr>
<tr><td>Separation column</td><td>12.5 cm × 75 µm capillary column<br>packed with 3 µm C18-silica particles<br>(Nikkyo Technos, Japan)</td></tr>
<tr><td>Flow rate</td><td>300 nL/min</td></tr>
<tr><td>Acetonitrile gradient<br>(in the presence of<br>0.1 % formic acid)</td><td>10 - 40%, 70 min</td></tr>
<tr><td rowspan="4">MS1</td><td>Mass resolution</td><td>70,000</td></tr>
<tr><td>AGC target</td><td>3.0E6</td></tr>
<tr><td>Maximum IT</td><td>60 msec</td></tr>
<tr><td>Scan range</td><td>310 - 1500 m/z</td></tr>
<tr><td rowspan="8">MS2</td><td>Mass resolution</td><td>17,500</td></tr>
<tr><td>AGC target</td><td>5.0E5</td></tr>
<tr><td>Maximum IT</td><td>60 msec</td></tr>
<tr><td>Loop count (Top N)</td><td>10</td></tr>
<tr><td>Isolation window</td><td>3.0 m/z</td></tr>
<tr><td>NCE</td><td>30</td></tr>
<tr><td>Dynamic exclusion</td><td>15 sec</td></tr>
<tr><td>Charge exclusion</td><td>1, 5-8, >8</td></tr>
</table>

**Supplementary Table 6.** Data collection, processing, model refinement and validation statistics.

| | PepNL-70S-RF2 | PepNL-70S |
|---|---|---|
| **Data collection and Processing** | | |
| Microscope | Titan Krios | Titan Krios |
| Voltage (kV) | 300 | 300 |
| Camera | Gatan K3 Camera | Gatan K3 Camera |
| Magnification | 105,000 | 105,000 |
| Pixel size at detector (Å/pixel) | 0.83 | 0.83 |
| Total electron exposure ($e^-$/Å$^2$) | 30 | 30 |
| Exposure rate ($e^-$/pixel/sec) | 14.3 | 14.3 |
| Number of frames | 30 | 30 |
| Defocus range (μm) | −0.8 to -1.6 | −0.8 to -1.6 |
| Energy filter slit width (V) | 20 | 20 |
| Micrographs collected (no.) | 15,342 | 15,342 |
| Final particles (no.) | 71,980 | 87,514 |
| Point group | $C_1$ | $C_1$ |
| Resolution (global, Å) FSC 0.143 (masked) | 2.90 | 2.85 |
| Map-sharpening $B$ factor (Å$^2$) | -58.2 | -65.8 |
| **Model composition** | | |
| Atoms | 143,036 (Hydrogens: 0) | 140202 (Hydrogens: 0) |
| Chains | 66 | 65 |
| RNA residues | 4461 | 4461 |
| Protein residues | 5971 | 5614 |
| Metal ions | 350 | 350 |
| Ligands | 8 | 8 |
| **Model Refinement** | | |
| Model-Map CC (mask/ box/ peaks/ volume) | 0.86/0.82/0.78/0.83 | 0.87/0.79/0.78/0.83 |
| Resolution (Å) by model-to-map FSC, threshold 0.50 (masked/ unmasked) | 3.0/3.0 | 2.9/2.9 |
| Average $B$ factor (Å$^2$) (RNA/ protein/ metal ion or ligand) | 30.73/35.20/25.73 | 21.65/25.09/16.72 |
| R.m.s. deviations, bond lengths (Å)/ bond angles (°) | 0.002/0.477 | 0.002/0.480 |
| **Validation** | | |
| MolProbity score | 1.37 | 1.35 |
| CaBLAM outliers (%) | 1.34 | 1.33 |
| Clash score | 6.71 | 6.32 |
| Rotamer outliers (%) | 0,08 | 0.41 |
| $C_\beta$ deviations | 0.00 | 0.00 |
| Ramachandran plot (%) (Favored/ allowed/ disallowed) | 98.43/1.57/0.00 | 98.64/1.36/0.00 |
